# Conserved heterodimeric GTPase Rbg1/Tma46 promotes efficient translation in eukaryotic cells

**DOI:** 10.1101/2020.07.06.190082

**Authors:** Fuxing Zeng, Xin Li, Melissa Pires-Alves, Xin Chen, Christopher W. Hawk, Hong Jin

## Abstract

Conserved developmentally-regulated GTP-binding (Drg) proteins and their binding partner Dfrp proteins are known to be important for embryonic development, cellular growth control, differentiation and proliferation. Here, we report that the yeast Drg1/Dfrp1 ortholog, Rbg1/Tma46, facilitates translational initiation, elongation and termination via suppressing prolonged ribosome pausing. Consistent with the genome-wide observations, Rbg1 reverses the growth defect resulting from translation stalling and stabilizes mRNAs against no-go decay. Furthermore, we provide a cryoEM structure of the 80S ribosome bound with Rbg1/Tma46 that reveals the molecular interactions responsible for Rbg1/Tma46 function. The Rbg1 subunit binds to the GTPase association center of the ribosome and the A-tRNA, and the N-terminal zinc finger domain of Tma46 subunit binds to the 40S, establishing an interaction critical for the complex ribosomal association. Our results answer the fundamental question on how a paused ribosome resumes translation and show that Drg1/Dfrp1 proteins play a critical role in ensuring orderly translation.

## INTRODUCTION

Maintaining protein homeostasis is essential for cell physiology, and this process, undoubtedly, is closely related to the accuracy and efficiency of protein synthesis. Disrupting intracellular homeostasis underlines a wide range of human diseases ^1–3^. The central translation apparatus, the ribosome, is a major target of control through its interactions with diverse proteins^4–6^. One of those proteins, the conserved <u>d</u>evelopmentally-regulated <u>G</u>TP-binding proteins (Drg), plays important roles in embryonic development, cellular growth, differentiation, and proliferation. Drg was originally identified to be highly expressed in neural precursor cells in the developing mouse brain ^7^. Shortly after its discovery, Drg mRNA and protein were found to be widely expressed at variable levels in cultured cells, as well as other embryonic, postnatal, and adult murine tissues ^8^. The coding sequence of Drg proteins contains a G-motif common to the GTPase superfamily ^9, 10^. A phylogenetic study revealed that eukaryotes typically contain two Drg genes, Drg1 and Drg2, that have highly homologues amino acid sequences (**Supplemental Figure 1**), whereas archaea contain only one ^11^.

In the cell, expression of Drg proteins is controlled by Drg family regulatory proteins (Dfrp) through direct physical associations: Dfrp1 specifically binds Drg1, whereas Dfrp2 binds to Drg2 preferentially ^12^. Like the Drgs, the sequences of Dfrp proteins are highly conserved. Yeast and mammalian Dfrp proteins share a partially-conserved sequence of about 60 amino acids, the DFRP domain, which is critical for binding to Drg proteins (**Supplemental Figure 1**). Importantly, association of Dfrp and Drg proteins confers stability to the Drg protein in vivo ^12^ and enhances the GTPase activity of the Drg in vitro ^13, 14^.

Both Drg and Dfrp proteins are conserved from yeast to human suggesting that they play important functions in fundamental pathways in eukaryotic cells. These proteins are highly expressed in actively growing and developing cells, as well as reproductive adult tissues of plants, animals, and humans. Consistent with their functions in growth control, altered Drg expression leads to cell transformation or cell cycle arrest ^15–18^. While high levels of Drg expression are positively correlated with their functions in translation, and both Drg1/Dfrp1 and Drg2/Dfrp2 complexes copurify with translation factors ^19^, the role played by these proteins in translation is not understood. Here, we report molecular functions of the yeast Drg1/Dfrp1 ortholog, Rbg1/Tma46, in translation and the structural basis of their mechanism of action on the ribosome.

## RESULTS

To effectively capture and compare *in vivo* ribosome dynamics in a drug-free way at a genome-wide scale, we employed the 5PSeq method (**Supplemental Figure 2A and 2B**), which captures 5’ monophosphate (5P) mRNA intermediates, produced by 5’ exonucleases (Xrn1 in yeast) that follow the last translating ribosome on an mRNA ^20^. As a result, 5PSeq enriches ribosome footprints on the mRNA undergoing 5’ to 3’ co-translational degradation, providing a sensitive measure of ribosome dynamics in translation and quality control pathways. Using this technique, we observe large a degree of ribosome pauses on mRNAs, even in the wild-type cell, showing that translation pause is likely to be ubiquitous in the cell.

In parallel, we created the Δrbg1 strain and multiple knockouts of genes functionally related to Δrbg1. The obviously reduced fitness of yeast cells as a result of double mutants of the Drg/Dfrp family members (Rbg1, Rbg2, Tma46, and Gir2) ^21^ suggests at least a partially-overlapping functional consequence between the two Drg/Dfrp complexes. Furthermore, yeast genetic screens followed by biochemical investigations showed that one of either the Rbg1/Tma46 or Rbg2/Gir2 complexes was required for uncompromised growth of cells lacking the slh1 gene ^19^. The protein product of this gene, Slh1, is another highly conserved eukaryotic protein that associates with ribosomes ^19^ and it is known to be important for the ribosome-associated quality control (RQC) ^22–26^. Simultaneous functional inactivation of Rbg1, Rbg2, and Slh1 results in serious growth defects ^19^. Therefore, we created a conditional triple-knockout strain, Δrbg2Δslh1-Rbg1d, by simultaneously inhibiting Rbg1 mRNA transcription and promoting Rbg1 protein degradation ^27, 28^ . This way, when combined with the single (Δrbg1) and double (Δrbg2Δslh1) knockouts, cellular functions of Rbg1 can be specifically examined and the partially overlapping functions of the three proteins, Rbg1, Rbg2, and Slh1, can also be studied (**Supplemental Figure 3A**).

Similar growth phenotype patterns were observed at 19, 30, and 37°C for the wild-type and all mutant strains (**Supplemental Figure 3B**), suggesting the conditional knockout strain Δrbg1Δslh1-Rbg1d works in the manner expected, and the cellular function of Rbg1 is temperature-independent. Furthermore, a progressive decrease of the amount of polysomes and increase in the amount of 40S, 60S, and 80S ribosomal fractions were observed when cells depleted these three proteins one by one, demonstrating a gradual decrease of global translation as cells are depleted of these three proteins one after another (**Supplemental Figure 3C**).

### 1. Deletion of Rbg1 leads to accumulation of ribosomes at the start and stop codons

Using the 5P-Seq method, metagene analyses demonstrated progressively increasing peaks at the -14 nt and 4 nt positions in Δrbg1, Δrbg2Δslh1 and Δrbg2Δslh1-Rbg1d cells (**Figure 1A ii-iv**, solid lines in blue) when compared to the randomly fragmented mRNA control (dotted lines in black) and the wild-type cells (**Figure 1A, i**). Since the length of ribosomal footprints from the ribosomal P-site to its protected 5’-end is 14 nt in this experiment, these two positions correspond to the ribosome pausing at the start codon and after seven amino acids have entered the exit tunnel, respectively (**Supplemental Figure 4A and 4B**).

**Figure 1.**
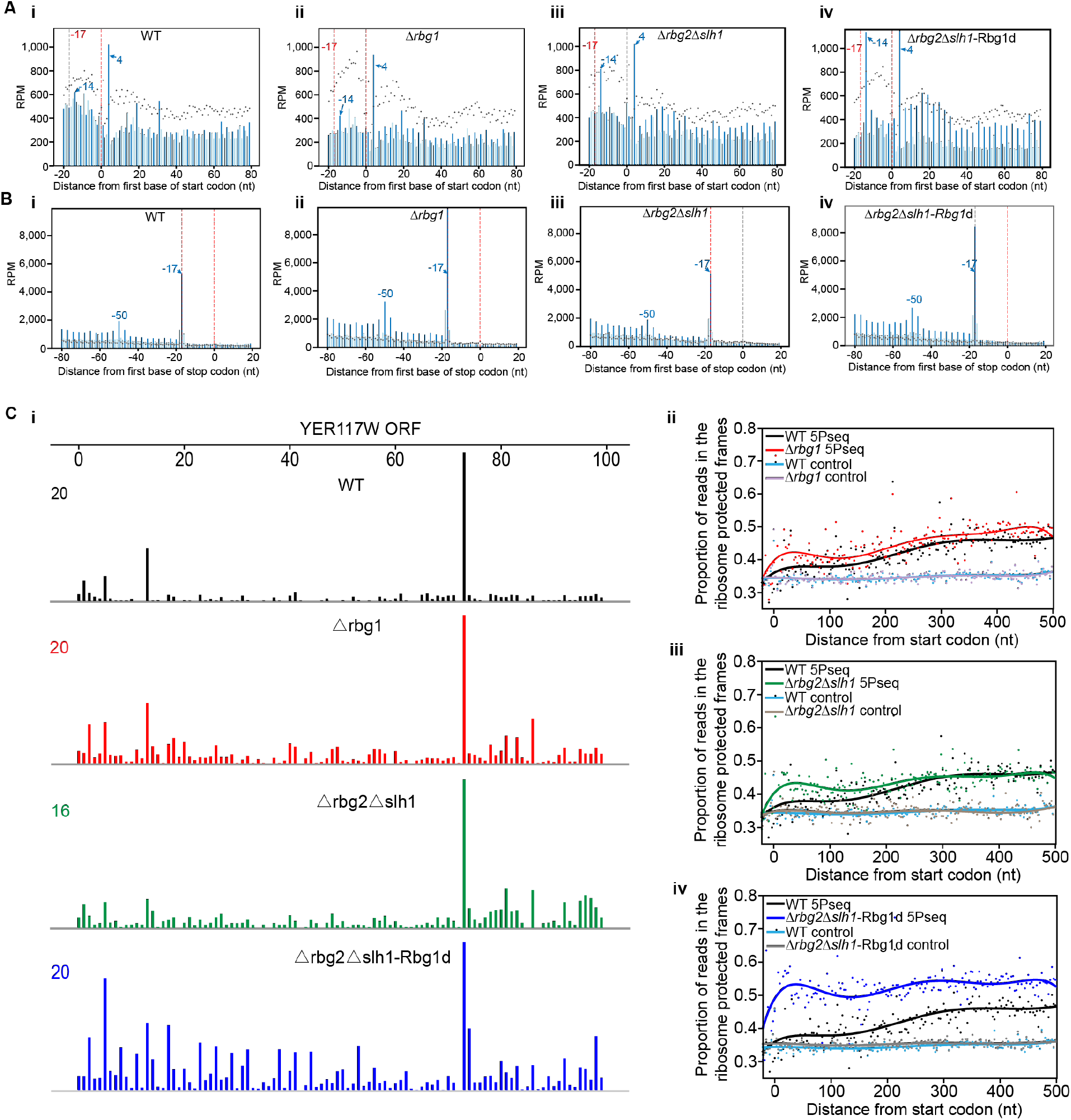
Deletion of Rbg1 results in slow translation initiation, elongation and termination. **A. Translation initiation arrests in mutant cells.** Normal translation pauses at start codons in wild type cells; and increased ribosome pauses at start codons in Δrbg1, Δrbg2Δslh1, and Δrbg2Δslh1-Rbg1d cells. Metagene analysis displaying the abundance of 5’P reads relative to start codons for WT, Δ*rbg1*, Δ*rbg2*Δ*slh1* and Δ*rbg2*Δ*slh1*-Rbg1d strains, or after random fragmentation (5PSeq control, dotted black line). Reads are represented by RPM, with the blue bar indicating the +1 frame and light blue bars indicating the 0 and +2 frames. The two dotted red lines show the positions of 0 and -17nt relative to the start codon. The blue peak at -14nt corresponds to the protected region from a putative initiation-paused ribosome and the peak at 4nt is caused by peptide-induced ribosomal arrest. Biological replicates are averaged. **B. Effects on translation termination in mutant cells.** Normal translation pauses at stop codons in wild type cells and altered pauses in the three mutants Δrbg1, Δrbg2Δslh1, and Δrbg2Δslh1-Rbg1d. Peaks at the -17nt position denote ribosomes with a stop codon in the A site; and the pause at the -50nt position is indicative of a disome position, with the leading ribosome reaching translation termination. Other experimental details are the same as described in **A**. **C. Slower translational elongation after depletion of Rbg1, Rbg2, and Slh1. i.** Representative genome tracks of the 5’ ends of 5PSeq reads in WT (black), Δ*rbg1* (red), Δ*rbg2*Δ*slh1* (green), and Δ*rbg2*Δ*slh1*-Rbg1d strains (blue). Coverage is expressed in RPM. An obvious 3-nt periodicity pattern was observed for YER177W mRNA in the Δrbg2Δslh1-Rbg1d strain. ii-iv: The proportion of 5PSeq reads in the ribosome-protected frame (Frame 1 in Supplementary Figure 2) shows that the 5’ end of mRNA was protected by ribosomes in the Δrbg2Δslh1-Rbg1d strain. Proportion scores were calculated for each codon by normalizing the read counts in frame 1 to the total reads in all three frames within the same codon. These are shown as dots with smoothed lines (polynomial fitting) for genes longer than 1000 base pairs (bp) and with RPKM values > 20. Only the region containing -20 nt to 501 nt with respect to the first base of start codon were used to calculate the proportion score. 5Pseq samples from Δ*rbg1* (ii, red), Δ*rbg2*Δ*slh1* (iii, green), and Δ*rbg2*Δ*slh1*-Rbg1d (iv, blue) strains were compared to WT strains (black). Random fragmentation samples (control) are also shown. Biological replicates are averaged.

At the stop codon, we also observe defects of slowed termination in Δrbg1 and Δrbg2Δslh1-Rbg1d cells (**Figure 1B** and **Supplemental Figure 4C** and **4D**). In-line with the function of Slh1, Δrbg2Δslh1 only shows a modest defect, suggesting that elongation stalling resolved by Slh1 likely alleviates ribosome accumulation at the stop codon. In addition, we found that ribosome pausing at the seventh amino acid was specifically associated with peptides containing a MSxxxxx pattern (**Supplemental Figure 4E**) ^20^. Other than this, no consensus sequences at the amino acid or nucleotide levels were found at either the start or stop codons, nor was any evidence found that specific cellular pathways are targeted by this phenomenon (data not shown). These results show that functions of Rbg1 and Rbg2 are not likely limited to a specialized pathway.

### 2. Inactivation of Rbg1, Rbg2 and Slh1 results in redistribution and accumulation of ribosomes in the 5’ end of mRNA

The mRNA degradation machinery closely follows the last translating ribosome in 5PSeq, where the translation rate of the last ribosome affects the amount of 5P intermediates and their positions in the coding region of mRNA. As a result, a 3-nt periodicity pattern of the 5P intermediates in the coding region (CDS) informs about ribosome dynamics, where a strong 3-nt periodicity is suggestive of slow translation, and weak 3-nt periodicity indicate fast (**Figure 1C i)**. It has been suggested that in wild type cells, it’s difficult for the exonuclease to catch up to the last translating ribosome near the start codon because decapping was rate-limiting ^20^. Compared to those in the wild type, in Δrbg1, and Δrbg2Δslh1 cells, we observe a much clearer 3-nt periodicity pattern of the 5P intermediates in the 5’ end of coding regions in Δrbg2Δslh1-Rbg1d cells (**Figure 1A and 1C-i**), suggesting that the rate of translation elongation was significantly decreased when the cells lost the functions of Rbg1, Rbg2, and Slh1 simultaneously.

To characterize this phenomenon more quantitatively, we calculated the proportion of 5P reads in the ribosome-protected frame for each codon. The proportion score for a control sample, lacking the characteristic 3-nt periodicity pattern, was estimated to be around 0.33. In agreement with an earlier observation ^20^, the proportion scores of 5P reads slowly increase from ∼0.33 at the start codon to ∼0.45 at the 300nt position in the ribosome-protected frame for the wild type cells (**Figure 1C ii-iv, black lines**). In contrast, the proportion values at the same position of the mRNA increase noticeably faster in the Δrbg1 (red line in **Figure 1C ii**) and Δrbg2Δslh1 cells (green line in **Figure 1C, iii**). However, this value increases to 0.45 immediately after the start codon in the Δrbg2Δslh1-Rbg1d cells (blue line in **Figure 1C iv**), indicative of a substantial decrease in the rate of elongation in these cells.

### 3. Rbg1 alleviates ribosome pausing at specific amino acids and mRNA regions

Seemingly stochastic yet pervasive ribosome pauses were seen at the individual gene level in our data. However, at the metagene level, we observed that ribosome pausing is correlated with the translation of glutamic acids, aspartic acids, arginine and lysine (**Figure 2A**), as well as serine and glycine but to a lesser degree. Considering that the length of ribosomal footprints from the ribosomal A, P, and E-sites to its protected 5’-end is 17nt, 14 nt, and 11nt in the 5P-Seq experiment (**Supplementary Figure 4A and 4C**), the observed pause in the ribosome at the metagene level likely occurs at the amino acid instead of nucleotide level.

**Figure 2.**
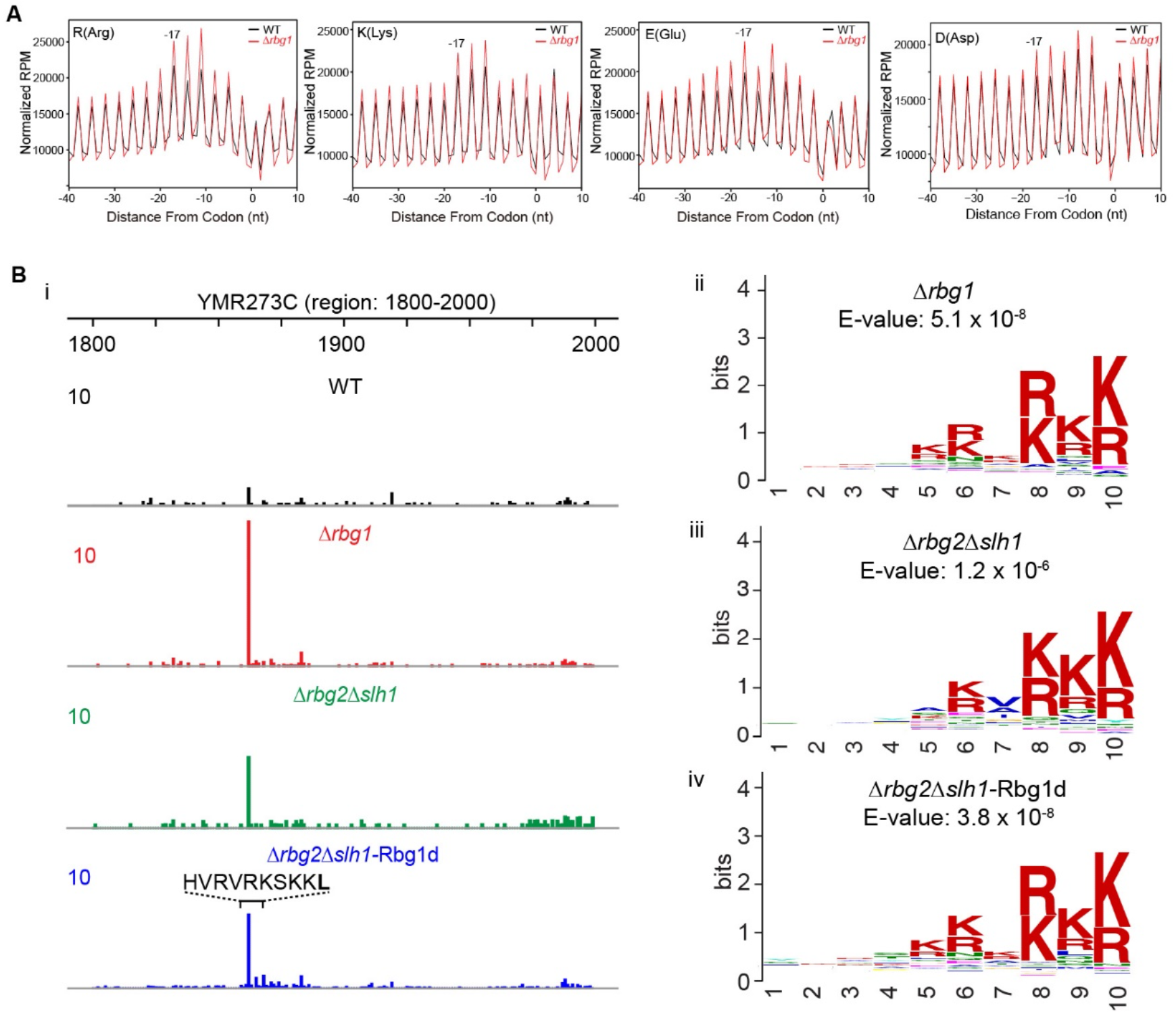
Rbg1 suppresses ribosome pause in translation. **A.** Increased translation pauses at Arg, Lys, Glu and Asp positions in the absence of Rbg1. The metagene (−40 to +10 window) which shows the number of 5P intermediates from translation of Arg, Lys, Glu and Asp amino acids and their ribosomal positions in *Δ*Rbg1 (red lines) and WT (black lines) cells. The total number of reads was normalized to facilitate data analysis and comparison, and data were analyzed as described ^33^. The -17nt, -14nt and -11nt peaks represent indicated codons at the ribosomal A, P and E sites, respectively. **B. Rbg1 alleviates ribosome pausing at arginine/lysine-rich regions. i**. **Representative genome tracks of 5’ ends of 5PSeq reads.** Genome tracks of the 5’ ends of 5PSeq reads in WT (black), Δ*rbg1* (red), Δ*rbg2*Δ*slh1* (green), and Δ*rbg2*Δ*slh1*-Rbg1d (blue) strains were shown around the arginine-/lysine-rich region. Coverage is expressed in average RPM of the biological duplicates. **ii-iv. Translation pauses at the R-/K-rich region in Δ*rbg1*, Δ*rbg2*Δ*slh1,* and Δ*rbg2*Δ*slh1*-Rbg1d strains.** To identify specific peptide sequences which induce ribosome pausing while being translated in the knockout strains, pause scores were first calculated by dividing the RPM value at each position of the transcript by the mean RPM value for the ten codons upstream and downstream of the same position. The ten amino acids upstream with a pause score > 10 were compared to those with pause scores < 10 using MEME ^51^. Amino acid sequences extracted from the WT strain show no consensus sequence.

In agreement with the results at the start and stop codons, no specific cellular pathway is targeted by Rbg1 proteins, which suggests that they play a general role in suppressing translation pause. Heterogeneity of the peptide sequences is the hallmark of the translation pause observed in our genomic data. Nevertheless, when we calculated the pause score at each position as described with modifications ^29^, R/K-rich sequences stood out as one common feature in all Δrbg1, Δrbg2Δslh1 and Δrbg2Δslh1-Rbg1d (**Figure 2B**). This result correlates well with the strong pausing effects of these two amino acids in the ribosome, and enrichment analysis was carried out on the most affected R/K-rich mRNA transcripts when Rbg1, Rbg2 and Slh1 are absent (**Supplemental Figure 5**).

The conclusions drawn from the genomic investigations described above were tested by comparing the rate at which wild type cells grow in the presence of anisomycin to the rate at which cells lacking Rbg1 grow under the same conditions. The antibiotic anisomycin is known to cause translational stalling by binding to the peptidyl transferase center (PTC) of the ribosome^30^. In the presence of 10 µg/ml anisomycin, the Rbg1 deletion strain grows slightly, but observably slower on YEPD plates than wild type yeast cells (**Figure 3A**, the second line of cell growth), but this growth defect is revered when Rbg1 proteins is expressed in Δrbg1 cells (**Figure 3A**, the third line of cell growth).

**Figure 3.**
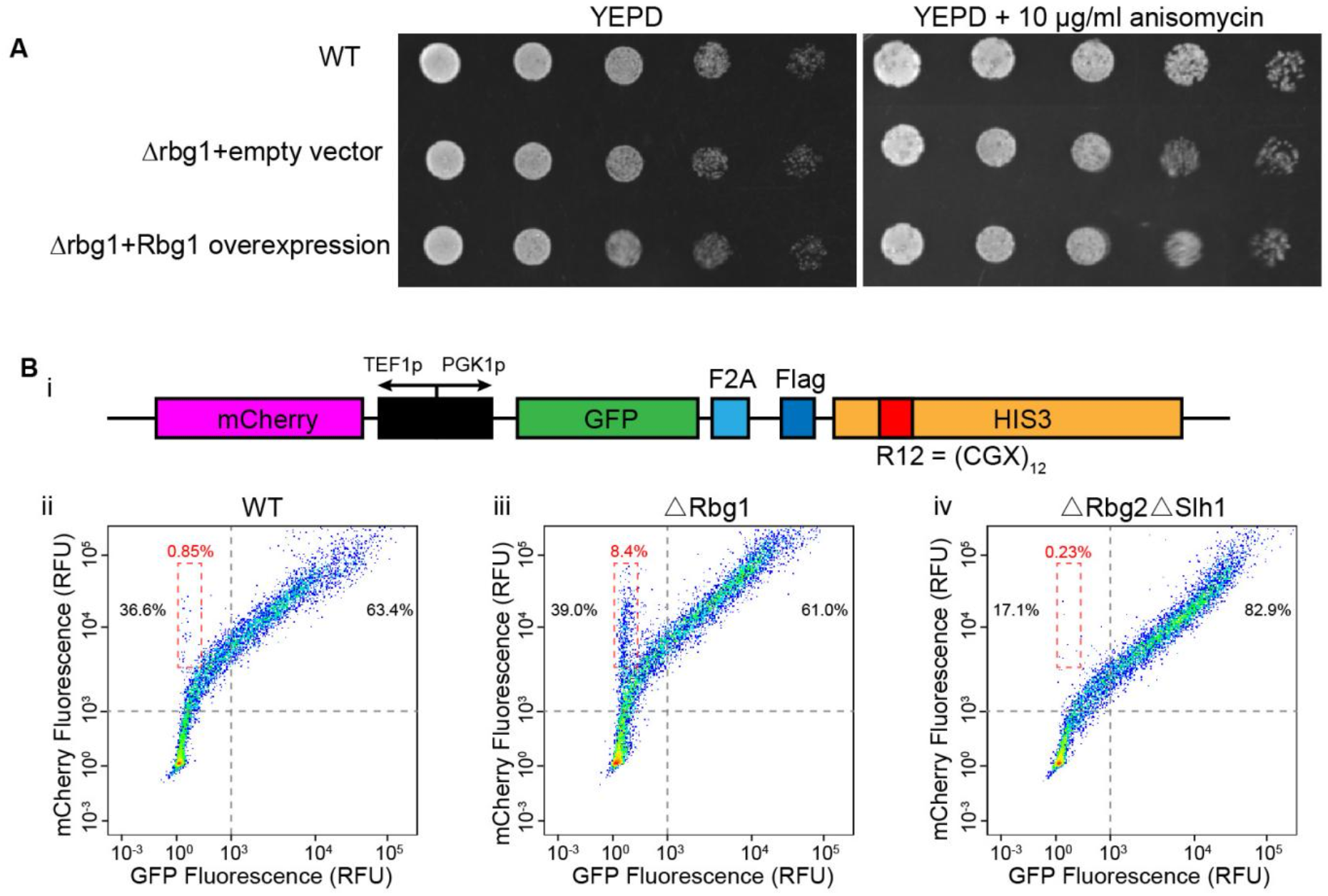
Rbg1 promotes efficient translation. **A. Deletion of Rbg1 results in slower growth for yeast cells in the presence of anisomycin.** Expression of Rbg1 protein in the Δrbg1 cells restored growth. Δrbg1 cells were transformed with either an empty vector, or a vector containing the rbg1 gene with a PGK1 promoter. Serial dilutions of exponential phase liquid cultures were spotted on YEPD, with or without 10 µg/ml anisomycin, and were incubated at 30 °C for 2 days. **B. Rbg1 enhances translation of mRNAs harboring an intrinsic stalling sequence.** Yeast cells expressing the indicated mRNA with the R12 sequence, fluorescent mCherry and GFP reporters (top panel) were grown to exponential phase (OD600=0.6) and were analyzed by flow cytometry. mCherry and GFP fluorescence intensities in the cell were monitored simultaneously using 561 nm and 488 nm excitation lasers, respectively. Scatterplots of 10,000 individual cells of the WT, Δrbg1 and Δrbg2Δslh1 background are displayed separately. The scatterplots are shown on a bi-exponential scale with pseudo-color in order to better visualize data across the wide range of expression levels seen in these experiments. According to the control cells which do not express the indicated mRNA construct (data not shown), only the cells with more than 10^3^ relative fluorescence unit (RFU) of mCherry fluorescence are considered to contain enough mRNAs and are subjected to further analysis. WT, Δrbg1 and Δrbg2Δslh1 cells with >10^3^ RFU mCherry intensities are divided into two groups, shown as percentages in grey dashed lines. Cells in the dashed box have high mCherry expression levels, but low GFP expression levels, the amount of which in WT, *Δ*Rbg1, *Δ*Rbg1*Δ*Slh1 is shown accordingly as percentages (red) (ii, iii and iv). The relative cell quantities at each position are indicated using a red to blue spectrum, where the red color represents the largest cell population, blue represents the smallest one.

To find out if Rbg1 facilitates the translation stalling on mRNA and protects mRNA harboring stalling signals from degradation that results from mRNA surveillance, we transformed cells with a construct mRNA containing twelve consecutive Arginine codons (R12), a sequence known to trigger mRNA no-go decay ^24, 31^, and monitor the degree of its translation using flow cytometry. Based on the experimental design (**Figure 3B, i)**, the intensity of the GFP fluorescence in counted cell populations is an indicator of the translation and the degree of stability of the construct mRNA. Compared to the WT cells (**Figure 3B, ii**), deletion of Rbg1 in the cell results in an increased number of cells with no GFP fluorescence (**Figure 3B, iii**), suggesting an increased degree of mRNA degradation. In contrast, the presence of Rbg1 in the Δrbg2Δslh1 cells leads to much enhanced GFP fluorescence, compared to not only the *Δ*Rbg1 cells, but also wild type cells. This result shows that Rbg1 indeed enhances the mRNA translation, stabilizes the mRNA and prevents them from undergoing no-go decay (**Figure 3B, iv**).

### 4. Structural basis of enhanced translation by heterodimeric GTPase Rbg1/Tma46 binding to the ribosome

To gain insights into molecular interactions important for the heterodimeric GTPase’s function, we obtained a cryoEM structure of the ribosome with Rbg1/Tma46 bound. A TAP tag was inserted at the end of the yeast chromosomal Rbg1 gene and the native Rbg1/Tma46- ribosomal complexes were purified from cellular ribosomal fractions using affinity pulldown. This method allows us to comprehensively sample physiologically-relevant functional states of the ribosome when the protein heterodimer binds. A more detailed experimental procedure is outlined in **Supplemental Figure 6**.

As shown in **Figure 4A**, we captured ribosomes in the classical ligand binding state with two tRNAs in the ribosomal A and P sites, protein eIF5A^28, 32, 33^ in the E-site and Rbg1/Tma46 bound to the GTPase association center (GAC). The ribosomes are in the translational state that they assume after peptide bond formation has occur but before tRNA translocation has taken place, with the nascent peptides attached to the CCA-end of the tRNA in the ribosomal A site.

**Figure 4.**
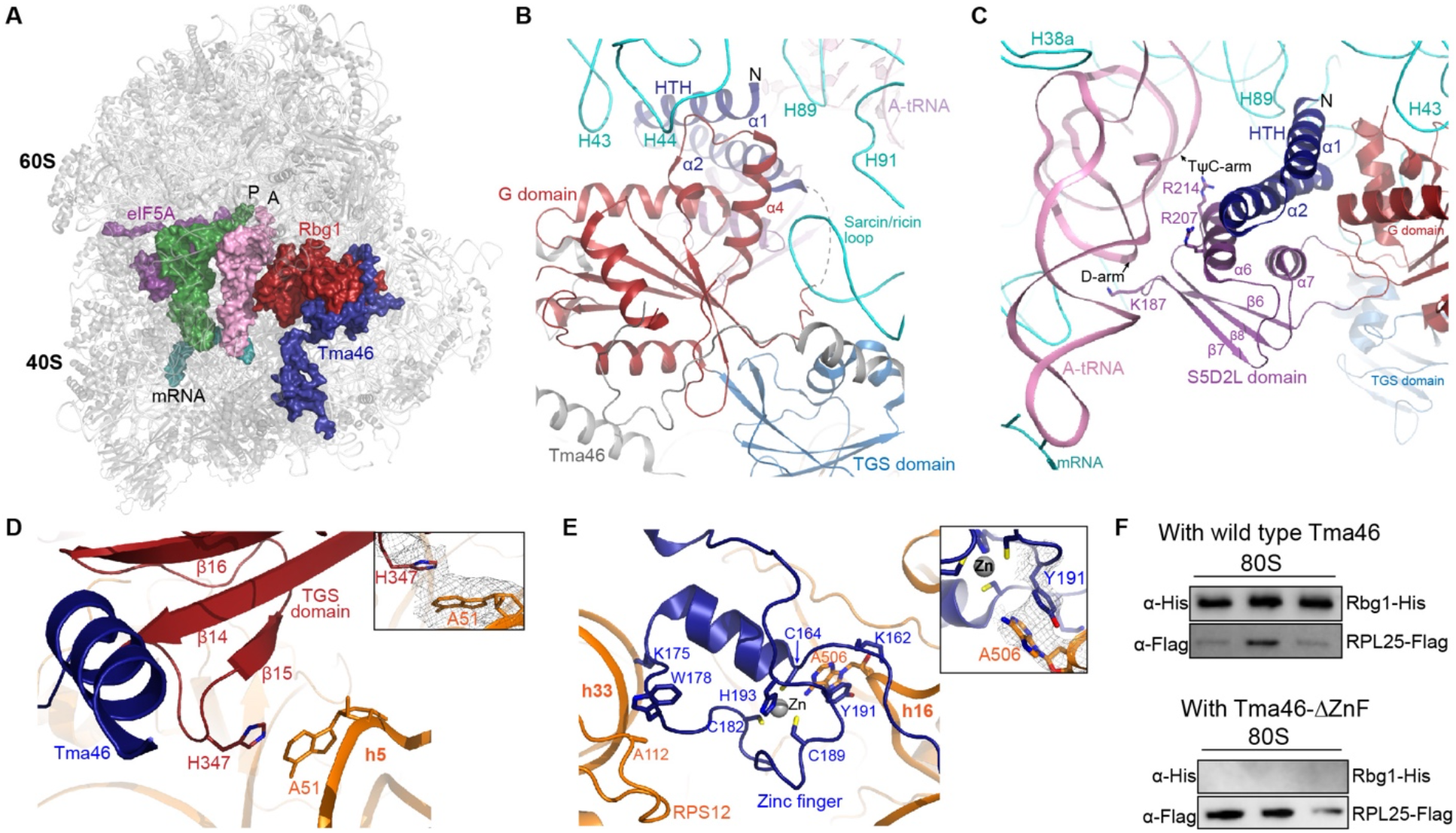
Structural basis for an enhanced translation when Rbg1/Tam46 binds to a paused ribosome. **A** An **o**verall structure of Rbg1/Tma46-bound ribosomes. The ribosome is colored in grey. P-tRNAs, A-tRNAs-nascent peptides, eIF5A, Rbg1 and Tma46 are colored in green, pink, magenta, red and blue, respectively. Detailed essential interactions of the Rbg1 G domain with the sarcin-ricin loop (**B**), S5D2L domain with A-tRNA (**C**), TGS domain with the h5 in the 40S subunit (**D**), and the second zinc finger in Tma46 with the head and shoulder regions of the 40S subunit (**E**). **F. The Rbg1/Tma46 complex binds to the ribosome via the zinc-finger domain of Tma46.** Proteins co-expressed in the *E.coli* BL21 strain were purified and incubated with pure 80S ribosomes, then ribosomal complexes were separated by sedimentation through a 10-50% sucrose gradient. Binding of Rbg1/Tma46 to the ribosome was determined by western blot. Flag-tagged RPL25 was immunoblotted on the same membrane. Both the GTP- and GDP-bound heterodimeric GTPases bound to the 80S ribosome, but only the GTP-bound state is shown.

The HTH (helix-turn-helix) and G domain of Rbg1 bind to the 60S subunit, with conserved residues in the *α*4 helix of Rbg1 interacting with the sarcin-ricin loop of the GAC (**Figure 4B**). Compared with the crystal structure of Rbg1/Tma46 (**Supplemental Figure 7**) in isolation, S5D2L (<u>S5</u> <u>D</u>omain 2 <u>L</u>ike) domain of Rbg1 extends out and charges residues in the loop regions between *β*6 and *β*7, as well as *α*6 and *β*8 interacts with A-tRNA (**Figure 4C**), and the TGS domain rotates ∼17 degrees and binds to h5 of the 18S rRNA (**Figure 4D**). These interactions are conserved (**Supplemental Figure 8**), suggesting that they are functionally important. Earlier studies showed that deletion of the TGS domain in Rbg1, or mutations in Rbg1’s G domain which either compromise the nucleotide-binding or GTPase activities of the protein, failed to rescue the severe growth defect in the triple knockout Δrbg1Δrbg2Δslh1 cell ^19^.

Tma46 adopts an extended structure and wraps around Rbg1. Their interaction is stabilized by extensive highly conserved hydrophobic contact. The second zinc-finger at the N-terminal region of Tma46 is ordered and binds to the gap formed between h16 and h33 of the 40S head and shoulder domains, respectively, establishing an interaction unique to the heterodimeric GTPase that is absent from other known monomeric translational GTPases such as EF-Tu, EF-G, RF3 (**Figure 4E**). This interaction is critical for mediating the ribosomal association of the heterodimeric GTPase. As shown in **Figure 4F**, deletion of the N-terminal Zinc-finger domain of Tma46 completely abolishes binding of Rbg1/Tma46 to the ribosome, regardless of whether Rbg1 is in the GDP or GTP-bound states. This result demonstrates that the zinc-finger domain of Tma46, while not required for Rbg1/Tma46 heterodimer formation ^13^, is required for ribosomal association.

## DISCUSSIONS

Translation proceeds at a non-uniform rate, with ribosomes pause as they progress down coding sequences on mRNA. As data demonstrated in this investigation, the ribosome pauses at every codon it encounters to execute the steps of the translation cycle, and that pausing is responsible for the 3-nt periodicity in the 5P-Seq. The pauses that occur at start and stop codons are longer than those seen in the elongation cycle because of the complexity of the events associated with initiation and termination. In the middle of coding sequences, variations in concentrations of cognate tRNAs as well as elongation factors play an important role in determining the rates at which particular codons are translated. Besides the “normal” pause, a prolonged pause of the ribosome could have effects ranging from “impediment but inconsequential” to detrimental, such as a translation arrest due to mRNA secondary structural elements, specific peptide motifs or damages in the mRNA ^34–38^. How ribosomes that have paused during the translation of a particular codon for a time longer than usual are recognized and targeted so that protein synthesis can continue is an important question. In this study, we show that in eukaryotes, conserved Drg proteins and their partners, Dfrps, play an important role in this process. The genomic experiments reported above show that the yeast ortholog of Drg1/Dfrp1, Rbg1/Tma46, reduces the pausing seen during initiation and termination, and enhances the overall rate of elongation. It is possible that the observed initiation arrest and elongation rate decline in all mutant strains, Δrbg1, Δrbg2Δslh1 and Δrbg2Δslh1-Rbg1d, are coupled. Remarkably, these negative impacts are accumulative, with the mildest defect occurring in the single knockout and the most severe defect occurring in the Δrbg2Δslh1-Rbg1d cell. Subsequent analysis indicates accumulated ribosome pauses underlines the observed phenomenon.

### Proposed Mechanism of Action

Our single-particle cryoEM reconstruction shows that the Rbg1/Tma46 complex targets translating ribosomes after peptide-bond formation has taken place, but before tRNA translocation. How does Rbg1/Tma46 promote efficient translation? In contrast to the altered conformations reported when stalling peptides reside in the exit tunnel without the Rbg1/Tma46 complex ^39–41^, the ribosomal decoding and peptidyl transferase centers assume conformations similar to what is observed in ribosomes competent to proceed further in the elongation cycle when Rbg1/Tma46 binds. This suggests that Rbg1/Tma46 acts by stabilizing pausing ribosomes in a productive conformation. Since this GTPase binds to the state immediately after peptide-bond formation, with the nascent peptide attached to the CCA-end of the A-tRNAs, it’s possible that binding of Rbg1/Tma46 facilitates tRNA interactions and peptide bond formation in the ribosome.

It is worth mentioning that the sheer number of ribosomal pause sites detected by the 5PSeq experiment, even in the wild-type cells, and the heterogeneity of the pausing sequences are striking to us, showing that translation pause is indeed a widespread phenomenon in eukaryotic cells. This phenomenon was also observed in several earlier studies ^28, 33, 42^ . Then*, how does Rbg1/Tma46 promote efficient translation in such a diverse molecular environment?*

Unlike other known monomeric translational GTPases, the heterodimeric GTPase Rbg1/Tma46 binds to the ribosome in both the GTP- and GDP-bound states in vitro (**Figure 4F**), indicating that it plays somewhat related, yet distinctive functions in translation. It’s reasonable to think that when translation pauses, E-tRNA leaves and vacates the ribosomal E-site. The zinc finger in the Tma46-subunit senses the movements in the 40S and binds to the gap between the head and shoulder domain. Stabilized by the Tma46-40S interaction, the nucleotide binding and GTP hydrolysis in the Rbg1-subunit might help the ribosome and tRNAs to navigate through a range of conformational and energetic landscapes resulted from heterogeneous pause signals, thereby achieving an efficient peptide-bond formation. Indeed, Nucleotide binding and GTP hydrolysis are important for both Rbg1’s association with the ribosome and its cellular functions^19^. In addition, Rbg1/Tma46 has a relatively low intrinsic nucleotide-binding and GTP-hydrolysis activity, this property may help the replacement of this GTPase by other elongation factors once their task is accomplished in the ribosome.

Because of the conserved nature of Rbg1/Tma46, a similar function is likely to be observed in higher eukaryotic organisms. Furthermore, the amino acid sequences of Drg1 and Drg2 are highly homologous with each other. Importantly, residues in Rbg1 that interact with the ribosome and tRNAs are either conserved or highly homologous, not only within the Drg1 family of proteins, but also between Drg1 and Drg2. This suggests that the Drg2 protein likely interacts with the translating ribosomes in a similar way and facilitates translation in a similar manner.

In contrast, amino acid sequences of Dfrp1 and Dfrp2, the Drg family regulatory proteins, are quite different except for their DFRP domain (**Supplemental Figure 1**). This suggests that the two Dfrp proteins are involved in different protein-protein interactions to engage Drg proteins to the ribosome, as also seen in yeast colliding ribosomes involve Gcn1 ^43^. Our structure explains earlier experimental observations that Dfrp1 specifically binds Drg1, and Dfrp2 binds to Drg2 preferentially ^12^. As seen in the structure, molecular interactions between Rbg1 and Tma46 are largely hydrophobic (**Supplemental Figure 1 and 8**). However, Rbg2 does not process the same amino acids with hydrophobic sidechains at the protein interface as Rbg1 does (**Supplemental Figure 8**), this molecular difference explains why Tma46 and Rbg2 does not bind together. On the other hand, some hydrophobic residues in Tma46 are conserved in Gir2, which has residues that could potentially interact with Rbg1 via other interactions at the protein interface, for example by formation of hydrogen bonds. Nevertheless, the weak cross-association of Drg1 and Dfrp2 ^44^ in the otherwise specific and distinct complexes, rules out the strict exclusiveness of the two Drg/Dfrp classes and suggests another fine-tuned layer of control in the cell.

Furthermore, while two Drg proteins likely play the same function on the ribosome, they are obviously involved in different translational events due to their distinctive Dfrp partners. Based on the data from flow cytometry (**Figure 3B**) where Rbg1 in the *Δ*Rbg2*Δ*Slh1 leads to stronger GFP signal produced from translation of mRNA with stalling sequences compared to the WT cells, Drg1 likely plays a more general role than Drg2 in the cell.

It came as a pleasant surprise that a general translation factor eIF5A was seen in the Rbg1/Tma46-bound ribosomal complex we studied. This protein is known to bind to the ribosomal E site ^45, 46^ and associates with the ribosome when translation stalls ^47, 48^. Translation at proline stretches is facilitated by eIF5A ^32^. While we indeed observe strong pause of ribosome when translating proline in wild type cells, deletion of Rbg1 does not exacerbate this effect (data not shown), suggesting mechanisms by which eIF5A and Rbg1/Tma46 promote translation likely to be different.

Translation pause can be viewed as a two-sided event: from the point of view of protein synthesis, a pause needs to be suppressed so that the protein synthesis can be resumed, and a protein can be made efficiently. However, on the other hand, a pause also provides a valuable chance for regulation. Whether Drg1/Dfrp1 acts as a suppressor of ribosome pausing at sites where mistakes (or suboptimal peptide bond formation) can occur is an important question for future study.

### Functional interplay of Drg1/Dfrp1, Drg2/Dfrp2 and ASCC3

The cumulative effects observed in this study among Δrbg1, Δrbg2Δslh1, and Δrbg2Δslh1-Rbg1d cells strongly suggest shared functions for Drg/Dfrp and ASCC3 (the mammalian counterpart of Slh1) proteins in fundamental biological processes. While it is still possible that the direct cellular targets and molecular functions of these proteins may not be the same, evidence available shows that the ribosome is one of their shared targets. Here, we propose these three proteins help to ensure cellular protein homeostasis by targeting ribosomes involved at the junction of protein synthesis and quality control pathways (**Figure 5**). When translation pauses, the Drg/Dfrp protein helps ribosomes to navigate through and find its way to pass the pause, thereby allowing the ribosomes behind it in the queue to continue using the mRNA as a template for protein synthesis. In the absence of Drg/Dfrp, ASCC3 likely guides the paused ribosome to the quality control pathway. Since the exact role of ASCC3 in the RQC or NGD is unclear, more investigations will be needed to reveal details at the molecular level.

**Figure 5.**
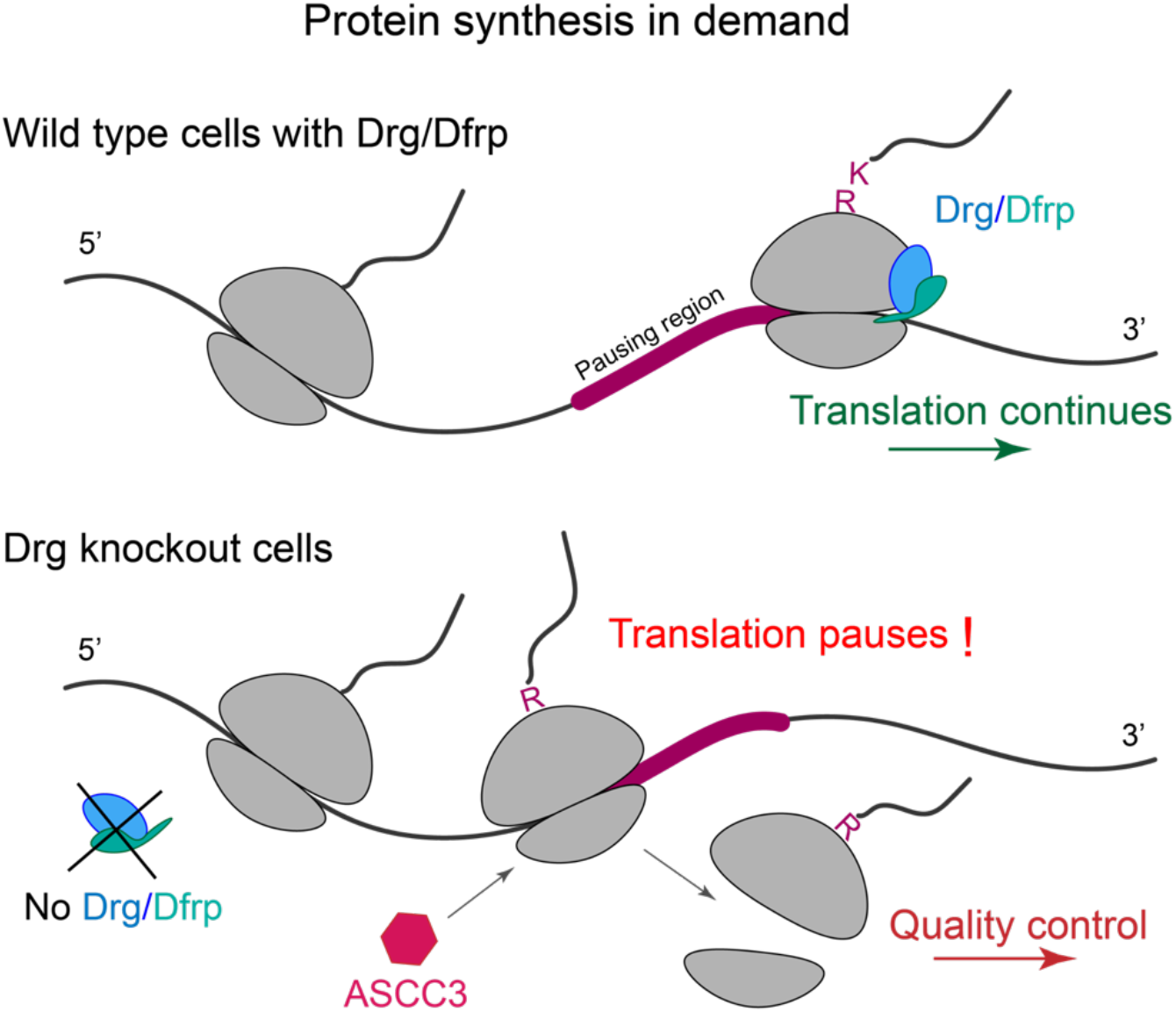
Functional interplay of Drg/Dfrp and ASCC in the translation pause-and-resume and quality control pathways. When translation pauses, Drg/Dfrp senses the paused ribosome and uses GTP-hydrolysis in the Drg subunit to promote efficient translation, whereby allowing subsequent ribosomes to continue translating the mRNA. In the absence of Drg/Dfrp, another protein called ASCC3, binds the stalled ribosome and triggers subunit disassociation, followed by mRNA degradation and no-go decay.

Whether Rbg1/Tma46 participates in ribosomal subunit disassociation is an important question to answer in future investigations. In addition, we found that it hydrolyzes ATP five times faster than GTP in vitro (Jin lab, unpublished data). It remains to be seen if association of this protein complex with the ribosome helps to couple metabolic and energetic state of the cell with efficient protein synthesis.

As a final note, Drg proteins play important roles in some cellular processes that appear to be unrelated to translation. For example, they are involved in spindle checkpoint signaling, and elevated levels of Drg1 protein causes lung adenocarcinoma and taxol resistance ^49^. In an *in vitro* study, Drg1 was reported to diffuse on microtubules, to promote microtubule polymerization, and to drive microtubule bundle formation in a GTP-independent manner ^50^. Additionally, HeLa cells with reduced Drg1 levels show delayed progression from prophase to anaphase due to a slowed spindle formation ^50^. Further investigations are required to see if these seemingly unrelated processes crosstalk with translation.

Maintaining protein homeostasis during critical stages and conditions such as development, proliferation, and cellular stress conditions is critical for eukaryotic organisms. It is not surprising that the translational machinery is one of the major targets of regulation for this purpose. Our observations provide links between translation pause and protein quality control, a critical component in maintaining protein homeostasis and cell physiology. It is worth mentioning that other pathways may crosstalk with these processes to achieve the same goal.

## METHODS

### Yeast strains, plasmid constructs and growth conditions

*Saccharomyces cerevisiae* strains with BY4742 (*MATα his3Δ1 leu2Δ0 lys2Δ0 ura3Δ0*) background were used in this study. Chromosomal insertion and knockout strains with this background were obtained by established homologous recombination techniques ^1, 2^, and are listed in **Supplemental Table 1**. The plasmids used here were constructed via standard cloning strategies and are listed in **Supplemental Table 2**. To generate C-terminal, chromosomally TAP-tagged strains, the TAP-*URA3* cassette, contained in the modified p415G plasmid, was amplified with homology regions to replace the stop codon of the gene to be tagged. Strains were confirmed by sequencing across the entire fusion gene, as well as western blot detection of the calmodulin binding peptide (CBP) epitope. It has been reported that cellular growth of the Δ*rbg1*Δ*rbg2*Δ*slh1*Δ strain was severely impaired ^3^. To control the Rbg1 protein level in the cell, a mini auxin-inducible degron (mAID) tag was inserted chromosomally into the Δ*rbg2*Δ*slh1* strain, at the 5’ end of the Rbg1 gene, with a sequence containing the *KanMX*-GAL10p-mAID-Flag epitope before the start codon of Rbg1 ^4^.

Genes encoding Rbg1 and Tma46 were amplified from the genomic DNA of S. cerevisiae and were cloned into modified pET28a or pET22b vectors using standard cloning strategies. Mutants were generated by mutagenesis PCR using the Q5 Site-Directed Mutagenesis Kit (NEB), according to the manufacturer’s instructions.

To obtain polysome profiles and perform 5Pseq experiments, wild type and mutant cells were grown to an OD600 of 0.6 in YPGR (2% galactose + 0.2% raffinose) medium at 30 °C, harvested by centrifugation, then washed and resuspended in the same volume of YEPD (2% glucose) medium containing 1 mM auxin (Sigma). These cells were grown in the presence of auxin for another 20 minutes at 30 °C to deplete Rbg1 completely.

### Purification of ribosomes, proteins

Yeast ribosomes from the YAS2488 strain were purified according to published methods^5^. Wild type and mutant Rbg1 and Tma46 were cloned and co-overexpressed in BL21(DE3) cells, then purified by affinity and ion-exchange chromatography ^6^.

### Ribosome binding assay

Purified Rbg1/Tma46 or Rbg1/Tam46-ΔZnF complexes were incubated with GDPCP or GDP for 15 minutes at 30 °C in polysome lysis buffer (20 mM Tris-HCl, pH 8.0, 140 mM KCl, 1.5 mM MgCl2, 1% (v/v) Triton X-100, 1 mM DTT, 100 µg/ml cycloheximide) followed by addition of purified 80S for another 30 minutes. The factor-bound ribosomal complexes were then separated by sedimentation through a 10-50% sucrose gradient in polysome lysis buffer. Binding of Rbg1/Tma46 to the ribosome was determined by western blot. Flag-tagged RPL25 was immunoblotted as control.

### Cell growth assay

Cell growth rates were determined by a spot assay as reported 7 with minor modifications. Cells were first grown to exponential phase at 30 °C in either YPGR or YEPD, depending upon the strain used. Then 0.5 OD600 units of cells were pelleted down and resuspended in 0.5 ml of 1X PBS buffer. The following serial dilutions were prepared in 1 x PBS buffer: 1/5, 1/25, 1/125, and 1/625, and 1 µl of each dilution was spotted on YEPD agar plates. 1 mM auxin (Sigma) was added for assays of Δrbg2Δslh1-Rbg1d strains. Images were taken after 2 to 5 days of growth at 30 °C.

### Polysome profiling

Yeast cells were grown at 30 °C until they reached an OD600 of 0.6, then were treated with 100 µg/ml of cycloheximide for 2 minutes before harvesting. Cells were quickly harvested by centrifugation and were washed twice with ice-cold polysome lysis buffer. The cell pellet was resuspended in 1 ml of polysome lysis buffer supplemented with EDTA-free protease inhibitors (Roche) and was grounded in liquid nitrogen using an RNase-free mortar and pestle. Extracts were clarified at 18,000 × *g* for 10 minutes at 4 °C, then loaded onto 10%-50% sucrose gradients in polysome gradient buffer (20 mM Tris-HCl, pH 8.0, 140 mM KCl, 5 mM MgCl2, 20 U/ml SUPERase.In, 0.5 mM DTT, 100 µg/ml cycloheximide). Gradients were centrifuged at 32,000 rpm for 3 hours and 45 minutes in a Beckman SW 32.1 Ti rotor and polysome profiles were generated by fractionation with continuous measurement of absorbance at 260 nm.

### Western blot

Western blotting was performed as described 8. Fractions from polysome profiles were resolved via SDS-PAGE, and proteins of interest were transferred to nitrocellulose membranes (GE Healthcare Amersham) for 40 minutes at 100 V in a Mini Trans-Blot apparatus (Bio-Rad). Protein bands were detected with either anti-Flag (Sigma-Aldrich) or anti-His antibodies (Sigma-Aldrich).

### 5Pseq library preparation

5Pseq libraries were prepared as described ^9, 10^. Yeast strains were grown in 50 ml YPGR medium until they reached an OD600 of 0.6. Cells were then harvested, and their media exchanged for 50 ml YEPD containing 1 mM auxin (Sigma), followed by an additional 20 minutes of incubation. Cells were harvested quickly by centrifugation. Total RNA was extracted, and DNA was removed by incubation with TURBO DNase (Ambion) for 30 minutes at 37 °C. For control libraries, 50 µg of DNA-free total RNA was fragmented by incubation at 80 °C for 5 minutes in RNA fragmentation buffer (40 mM Tris-acetate, pH 8.1, 100 mM KOAc, 30 mM Mg(OAc)2). Fragmented 5’-OH sites were phosphorylated by treatment with 5 units of T4 polynucleotide kinase (NEB) for 1 hour at 37 °C, followed by phenol:chloroform extraction. The phosphorylated 5’-ends were subjected to RNA ligation by 20 units of T4 RNA ligase 1 (NEB) in the presence of 10 mM DNA/RNA rP5_RND oligo (**Supplemental Table 3**) at 16 °C overnight. For the 5Pseq libraries, 50 µg of DNA-free total RNA was directly ligated to the rP5_RND oligo in the same conditions as specified for control samples. The polyadenylated mRNAs were enriched using Dynabeads (dT)25 (Life Technologies), according to the manufacturer’s instructions, followed by fragmentation for 5 minutes at 80°C in RNA fragmentation buffer. Both the controls and 5’ RNA-seq samples were primed with random hexamers and reverse transcribed with Superscript II (Life Technologies). Second strand cDNA synthesis was performed using a single PCR cycle (98 °C for 1 minute, 50 °C for 2 minutes, and 72 °C for 15 minutes) in Phusion High-Fidelity PCR Master Mix and was primed with BioNotI- P5-PET (**Supplemental Table 3**). Double-stranded cDNA was purified using HighPrep beads (Magbio), then was bound to Dynabeads M-280 Streptavidin beads (Life Technologies). Bound DNA molecules were subjected to end repair, adenine addition, and adaptor ligation via addition of 0.5 µL of P7MPX annealed adaptor, which was prepared by annealing the P7MPX_linker_for and P7MPX_linker_rev (**Supplemental Table 3**) primers at a final concentration of 2.5 µM. The Dynabead-bound DNA was washed and subjected to PCR amplification (98 °C for 30 seconds; 18 cycles of the following condition: 98 °C for 20 seconds, 65 °C for 30 seconds, 72 °C for 30 seconds; 72 °C for 5 minutes) using Phusion High-Fidelity PCR Master Mix with HF buffer (NEB), 0.1 µM of PE1.0, and the appropriate PE2_MPX_01 to PE2_MPX_08 primers (**Supplemental Table 3**). Samples were size-selected using 0.6x-0.9x (v/v) HighPrep beads (Magbio), separated via agarose gel electrophoresis, and the 300-500 bp regions of the gel were extracted. Libraries were sequenced using single-end, 100bp-read NovaSeq Illumina sequencing with 6 nt indexing reads for library identification.

### High throughput sequencing data processing and analysis

For all libraries, reads were first demultiplexed using the index sequences. The 3’ adaptor sequence was identified and removed using the cutadapt package ^11^. After adaptor removal, the remaining reads were deduplicated by bbmap (BBMap - Bushnell B. - sourceforge.net/projects/bbmap/) based upon random barcode sequences, and the 5’ UMI were trimmed by fastx_trimmer (Hannon, G.J. (2010) FASTX-Toolkit.). Non-coding RNAs were filtered by mapping to annotated S. cerevisiae rRNAs, tRNAs, snRNAs, and snoRNAs using Bowtie1 ^12^. Unaligned reads were then mapped to the sacCer3 genome using Bowtie2 13 with the arguments: ‘--local -D 15 -R 2 -N 1 -L 20 -i S,1,0.75 -S’.

Unless indicated otherwise, data analysis was performed using a series of custom scripts written in Python. For 5Pseq experiment, reads with multiple alignments, mapping quality values < 30, or those containing soft-clipped bases on the 5’ end were excluded from downstream analysis. The 5’ ends of reads passing quality filtering were extracted and the total numbers of reads per genome location were calculated. Reads per million (RPM) values for each genomic locus were calculated using the number of unique reads. Only genes with reads per million per kilobase (RPKM) values of greater than 20 were chosen for further analysis. To create meta gene plots, we calculated the average rpm of the 5’ ends from all biological replicates. These 5’ ends were then aligned relative to the stop codon (or any specific codon) of all ORFs, and the sum of the 5’ ends reads was determined at each position. To analyze ribosome pausing around the start codon, genes were sorted by a ratio value which is calculated by dividing the read counts at -14 nt or 4 nt by the total read counts corresponding to the surrounding ±2 codons, as previously described ^10^. Then the first 8 amino acids of the ORF were extracted. Sequences of the top 50% of genes were compared to the bottom 50% using MEME ^14^. To characterize the elongation pausing, we calculated the pause score, as described previously, with minor modifications ^15^. Pause scores were calculated by dividing the rpm value at each genomic position by the mean of the rpm values in the ±10-codon region surrounding the position. Regions in the first and last 10 codons of the ORF were excluded. Amino acid sequences with pause scores > 10 were considered to be stalling sites. Stalling sequences were calculated by MEME ^14^ using the sequences with pause score <10 as a control.

### Flow cytometry

Overnight cultured cells carrying the dual fluorescence reporter gene with a stalling sequence containing twelve consecutive arginine codons were first inoculated into 50 ml YEPD with initial OD600 = 0.1 and grown to mid-log phase at 30 °C. Cells were then collected and diluted to OD600=0.1 and about 10,000 cells were analyzed with a FACSCanto SORP flow cytometer (Made, company) for GFP and mCherry fluorescence detection using 488 nm and 561 nm excitation lasers, respectively. Scatter plots show the intensity of GFP and mCherry for individual cells. Flow cytometry was done in triplicate.

### Electron microscopy, data collection and image processing

Saccharomyces cerevisiae with C-terminally TAP-tagged Rbg1 were grown to log phase at 30 °C, then were treated with 100 µg/ml of cycloheximide for 2 minutes before being harvested by centrifugation. The harvested cells were resuspended in one-third of polysome lysis buffer and were grounded in liquid nitrogen. Cell extracts were clarified at 20,000 × *g* for 30 minutes at 4 °C and incubated with IgG beads for 1 hour at 4°C. The complex-bound beads were washed three times with polysome lysis buffer, and the Rbg1/Tma46 bound ribosomal complexes were eluted by incubating with TEV for 1 hour at 20 °C. 0.05% glutaraldehyde was added to the eluted complex and incubated for 30 minutes on ice. The crosslinking reaction was quenched by 25 mM Tris-HCl (pH 7.5) and 0.01% n-dodecyl-D-maltoside (DDM) was added to the final solution. Sample preparation for cryoEM studies was performed as described ^16^. 2.5 µl aliquots of ribosomal complexes were incubated for 30 seconds on glow-discharged Holley carbon grids with a thin-layer carbon film cover (Quantifoil). Grids were blotted using a Vitrobot Mark IV (FEI) for 3 seconds in 100% humidity at 4 °C, then were plunge frozen into liquid ethane. Data were collected in vitreous ice using Titan Krios G3i (D3796) transmission electron microscopes operating at 300 keV with FEI Falcon III detectors. A total of 11,529 micrographs were acquired using a dose of 30 e^-^Å^-2^. The drifts of movie frames were corrected using MotionCor2 ^17^, and the contrast transfer functions were determined using CTFFIND4 ^18^. Data processing was carried out in Relion3 ^19^. A total of 1,222,865 particles were picked and extracted for reference-free 2D classification. 737,448 particles were selected and subjected to 3D refinement program followed by 3D classification with a mask around the ribosome GTPase association center. The class displaying good factor density (95,380 particles) were subjected to 3D refinement to yield a reconstruction at 3.23 Å. Resolutions were reported based upon the gold-standard Fourier shell correlation (FSC) of 0.143 criterion ^20^.

### Model building, refinement and validation

The high-resolution crystal structure of the yeast 80S (PDB: 4V88) ^21^ was used and fit into the EM density map using rigid body fitting in Chimera^22^. The body, head and shoulder domains of 40S, L1 stalk and P stalk of 60S were fit separately using Coot 23. Ribosome-bound ligands, A-tRNA, P-tRNA and eIF5A, were built into density map in Coot using PDB 5GAK as a reference 24. The crystal structure of Rbg1/Tma46 complex (PDB: 4A9A) ^6^ was used a reference, and the HTH, S5D2L, G domain, TGS domain of Rbg1 and each helix of Tma46 were built into the density map accordingly. The second zinc finger of Tma46 was built using the homology model generated by SWISS-MODEL ^25^.

The model obtained was refined using Phenix with secondary structure, RNA base-pair, sugar pucker and base stacking restraints. The final model was validated using MolProbity 26. Refinement statistics for the structures were summarized in Supplemental Table 4. Figures were made in Chimera 22 and Pymol 27.

### Data availability

The sequencing data for 5P-seq experiment have been deposited in NCBI’s Gene Expression Omnibus and are accessible through GEO series accession numbers GSE154212. Electron microscopy maps have been deposited in the Electron Microscopy Data Bank under accession codes EMD-xxxx. Coordinates have been deposited in the Protein Data Bank under accession code xxxx.

## Acknowledgments

We thank Prof. Peter Moore at Yale University for critical comments of the manuscript. We also thank the Cryo-EM Center at Southern University of Science and Technology, Junyi Jiang and Qingrong Li in the F. Zeng laboratory, Dr. Bridget Carragher and Dr. Ed Eng at the National Center for CryoEM Access and Training (NCCAT) and Dr. Valerie Tokars at Northwestern University for their help on cryoEM data collection, the core Research Facility at Southern University of Science and Technology for flow cytometry, Roy J. Carver Biotechnology Center at the University of Illinois at Urbana-Champaign for sequencing and members in the H. Jin laboratory for helpful discussions. H.J. acknowledges support from the National Institute of General Medical Sciences of the NIH (R01-GM120552).

All authors declare no conflict of interests

## Supplementary Figures

**Supplemental Figure 1.**
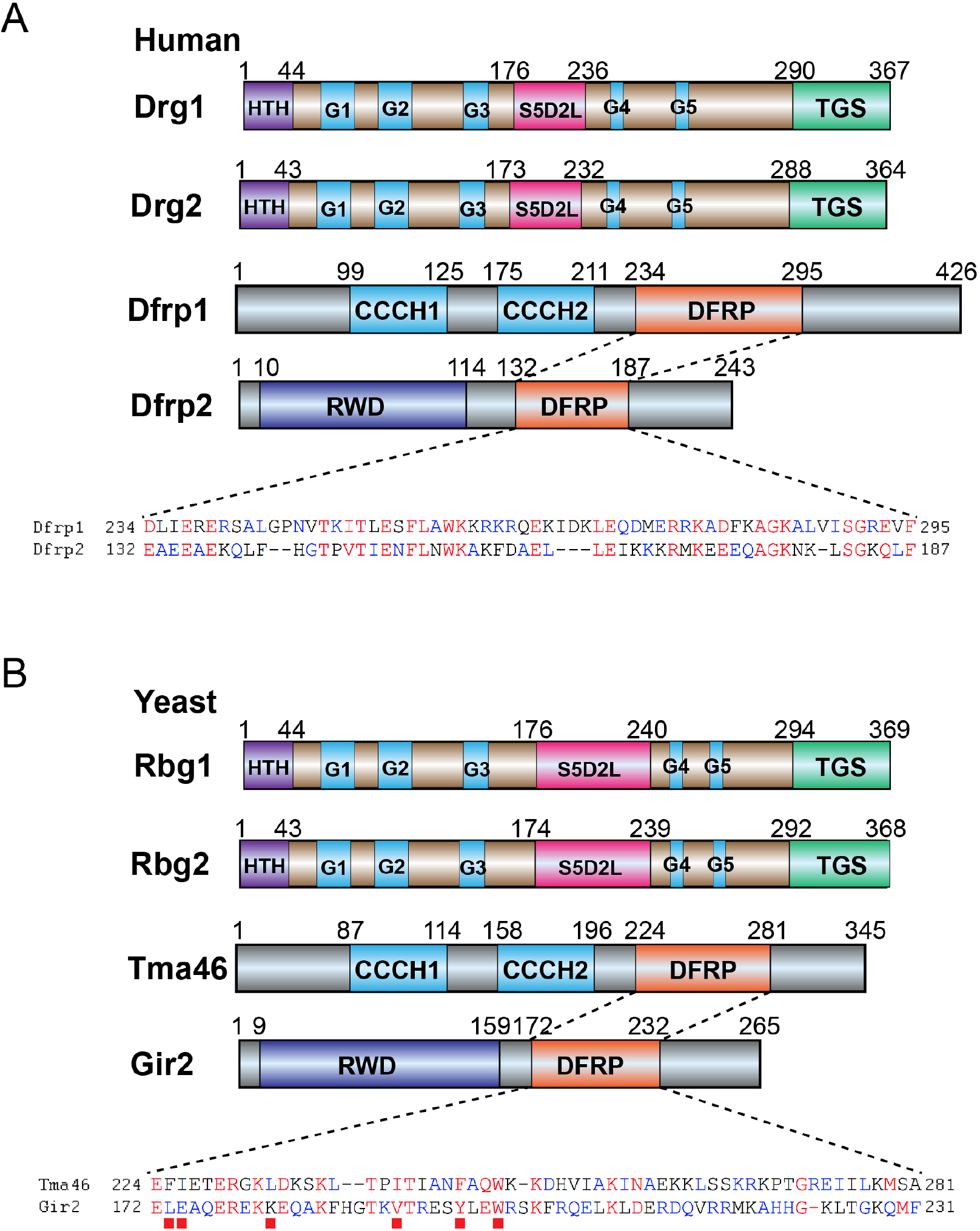
Overview of the domain organization of the Drg/Dfrp proteins in humans and yeast. Sequence alignments are shown for the DFRP domains. Red squares 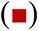 indicate hydrophobic residues in Tma46 that interact with Rbg1 when the heterodimeric complex Rbg1/Tma46 binds to the ribosome seen from the structure obtained in this study.

**Supplemental Figure 2.**
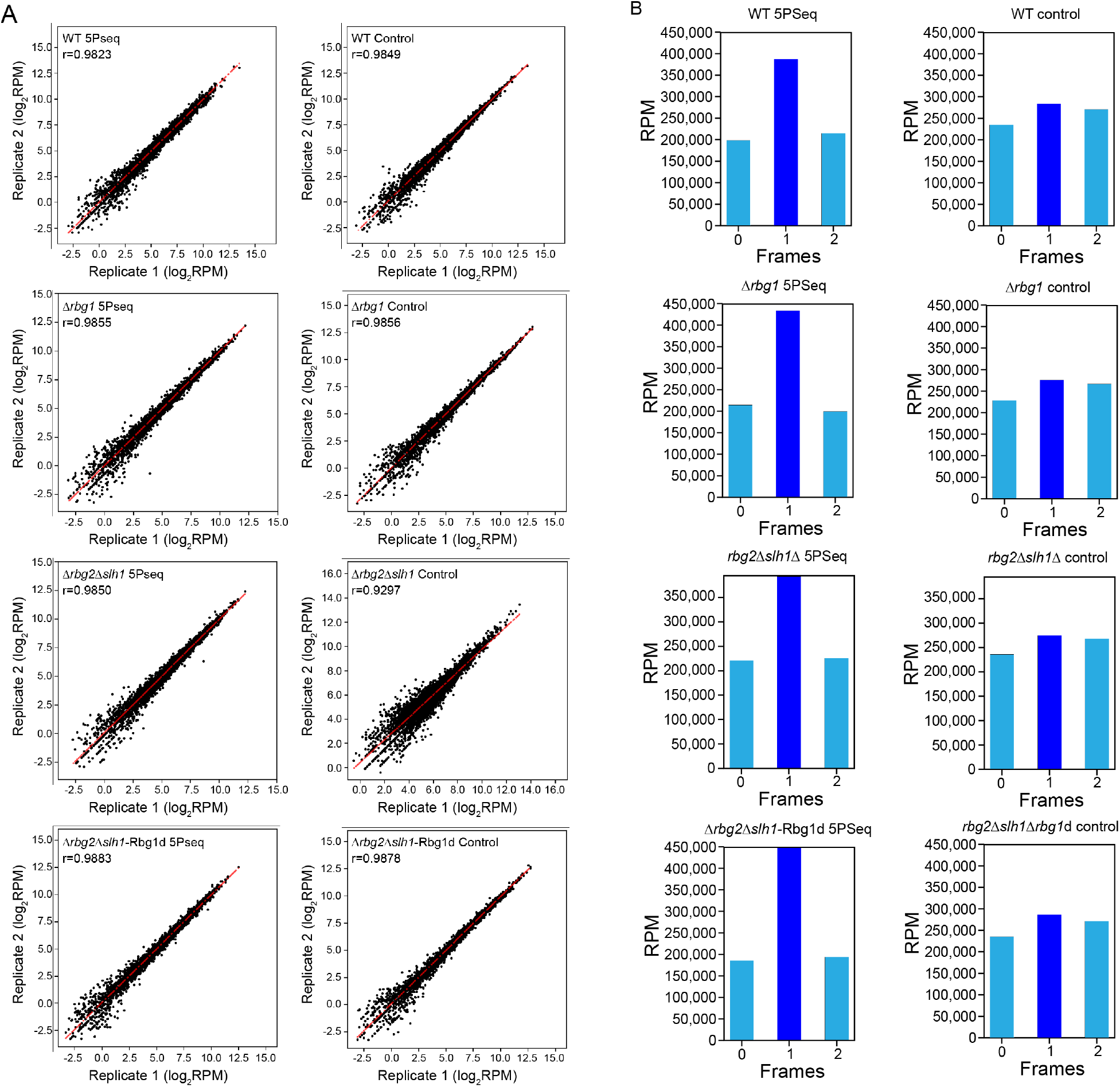
Data quality of 5PSeq experiments performed with different strains. **A.** Scatter plots showing reproducibility of 5Pseq data between biological replicates for 5Pseq (Left) and controls (Right). Reads with multiple alignments, mapping quality < 30, or those containing soft-clipped bases on the 5’ end were excluded. Reads were normalized to the library size using RPM (reads per million). Results for the two biological replicates are compared, and Pearson correlation coefficients for each comparison are shown. **B** 3-nt periodicity pattern of the 5P-seq data. The 5’ ends of reads were extracted and the total number of reads at each frame were calculated and are displayed as histograms for all of the codons along the coding region of all genes. Because the ribosome protected region in 5P-seq is 14 nt from the 5’ end to the P-site of the ribosome, the 5Pseq shows a peak at frame 1. Control samples were also analyzed in the same manner and show no preferred peak among the three frames.

**Supplemental Figure 3.**
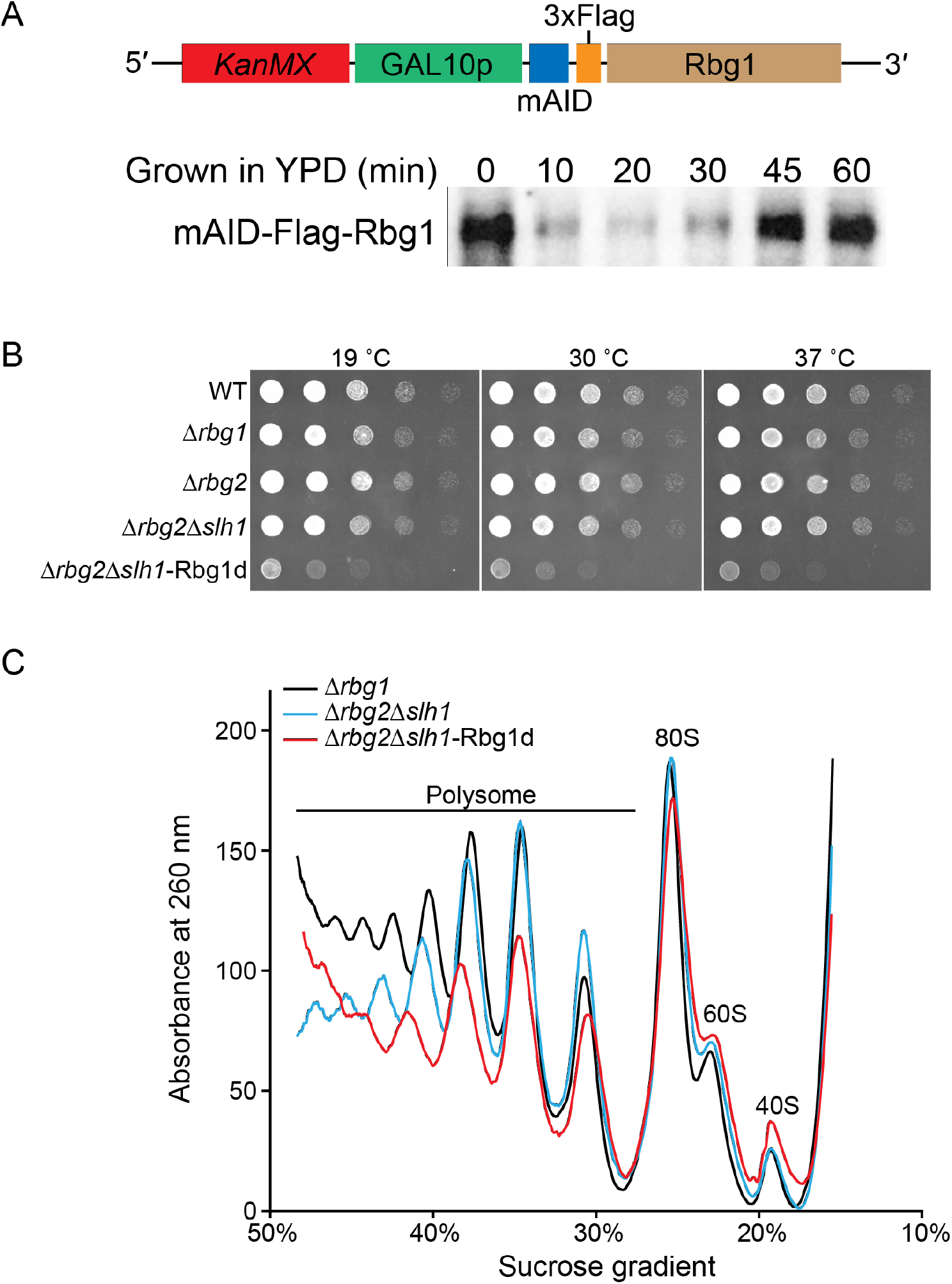
Rbg proteins play important roles in global translation. **A.** Model of the auxin inducible-degron system. To remove the Rbg1 protein from the cell, a GAL10 promoter was inserted into the 5’UTR of rbg1 gene so that the Rbg1 transcription levels can be controlled by the presence of glucose in the media. A mini auxin-inducible degron (mAID) tag was fused to the N-terminus of Rbg1. Cells were first grown in YPGR medium and then changed to YEPD medium containing 1 mM auxin to induce the degradation of Rbg1 protein in the cell. Aliquots at different time points were acquired and the Rbg1 protein levels were monitored by western blot with anti-Flag antibody. With the mAID degron tag, Rbg1 protein degrades after 20 minutes but its expression is restored after 45 minutes. **B**. Elimination of Rbg1 under Δrbg2Δslh1 background shows a temperature-independent growth defect. Serial dilutions of liquid cultures growing in exponential phase were spotted on YEPD plates containing 1 mM auxin and were incubated for 2 days at 30 °C or 37 °C, or for 4 days at 19 °C. **C**. Polysome profiles from cells incubated for 20 minutes in YEPD indicate that Rbg1 plays an important role in global translation. Cells were first grown in YPGR medium to OD600 0.6, then were changed to YEPD medium containing 1 mM auxin for 20 minutes. Cells were then quickly harvested and polysome profiling was performed. The profiles of WT and Δrbg2 strains are not shown here since WT, Δrbg2, and Δrbg1 have nearly identical profiles. These data represent one of three biological replicates.

**Supplemental Figure 4.**
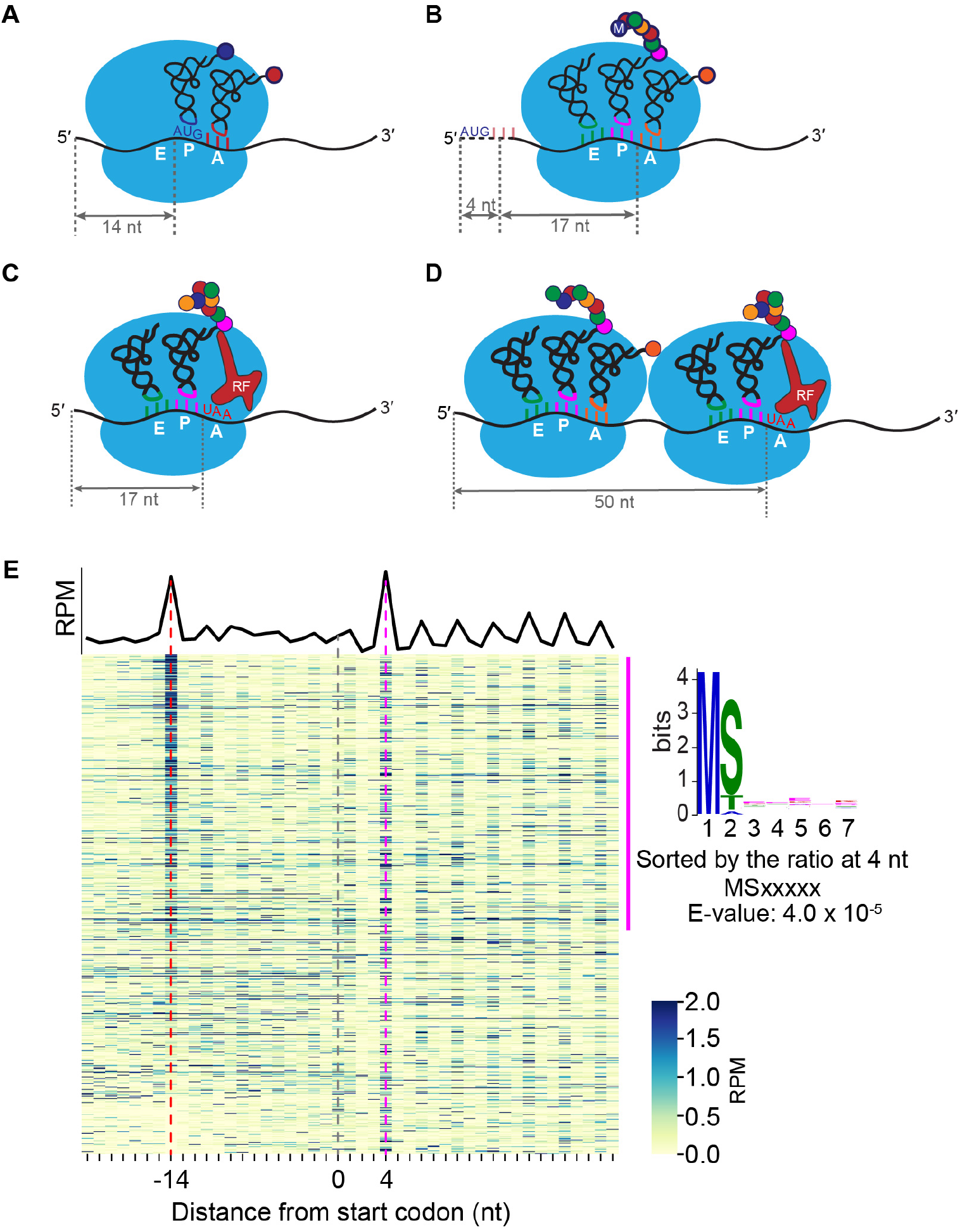
Ribosome pausing at the initiation and termination sites are sequence independent. **A to D**. Models for mRNA footprints generated by 5Pseq for stalling peaks at -14 nt (A) and 4 nt **(B)** at the initiation codon, and -17 nt (C) and -50 nt **(D)** at the termination codon. The length between the 5’ end and P-site in the traditional ribosome profiling method is 12-13 nt. Due to the steric hindrance between the ribosome and exonuclease Xrn1, the length between the 5’ end of the mRNA and the P-site is 14 nt for 5Pseq. **E.** A metagene representing both the average coverage (black line), and gene-specific coverage (blue heatmap) for the Δrbg2Δslh1-Rbg1d mutant. Only genes with RPM values larger than five in the displayed regions were considered. Genes were sorted by the ratio of reads at either the -14 nt (red dotted line) or 4 nt (magenta dotted line) positions to the reads corresponding to the surrounding ± 2 codons. To identify specific peptide sequences, the first eight amino acids of the genes in the top 50% were compared to the bottom half using MEME 1. No sequence motif was found by MEME for the genes sorted at -14 nt, and the sequence motif MSxxxxx was found for the genes sorted at 4nt position.

**Supplemental Figure 5.**
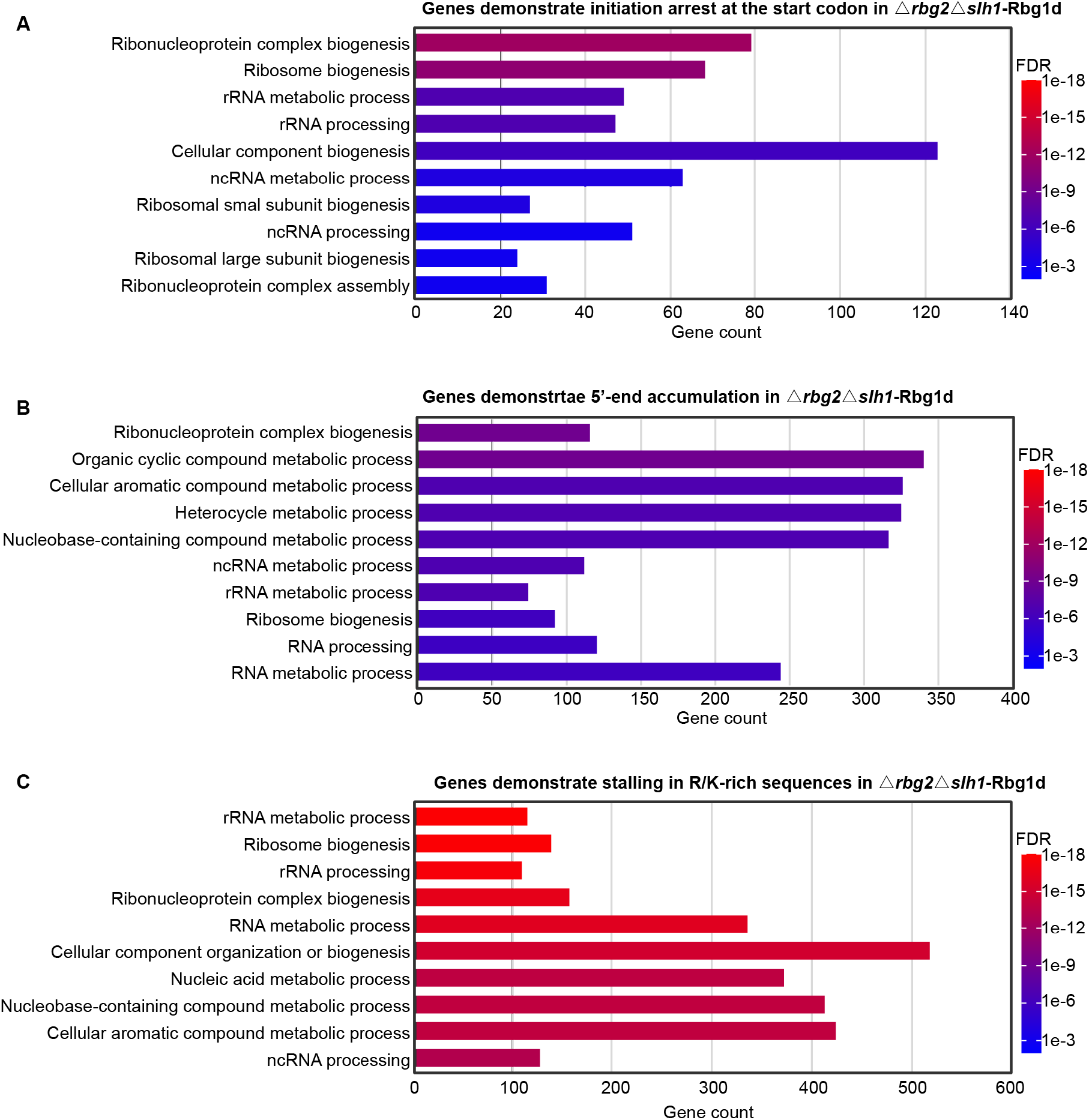
Gene ontology analysis for genes demonstrate initiation arrest, 5’- end accumulation and elongation stalling in R/K-rich sequences. **A.** Gene ontology analysis of genes showing ribosome pausing at the initiation site in the Δrbg2Δslh1-Rbg1d cell. By filtering the ratio of reads at -14 nt to the surrounding ± 2 codons larger than 0.1, 353 genes were selected from the Δrbg2Δslh1-Rbg1d strain for Gene ontology analysis using DAVID 2. The top 10 gene ontology terms for biological processes were shown, ranked by false discovery rate (FDR). **B**. Gene ontology analysis of the 715 genes which have higher proportion scores in the Δrbg2Δslh1-Rbg1d cell comparing to WT cell. The range of -12 nt to 99 nt, with respect to the first base of start codon, was used to calculate the proportion score. Genes whose proportion scores were larger than 0.4 in the Δrbg2Δslh1-Rbg1d cell, and whose proportion scores were more than 1.5 times the scores found in the WT cell were selected and analyzed by DAVID 2. The top 10 GO terms for biological processes are shown, ranked by false discovery rate (FDR). **C**. Gene ontology analysis of the 993 genes that have pause score larger than 10 in Δrbg2Δslh1- Rbg1d cell. The top 10 gene ontology terms for biological process were shown, ranked by false discovery rate (FDR).

**Supplemental Figure 6.**
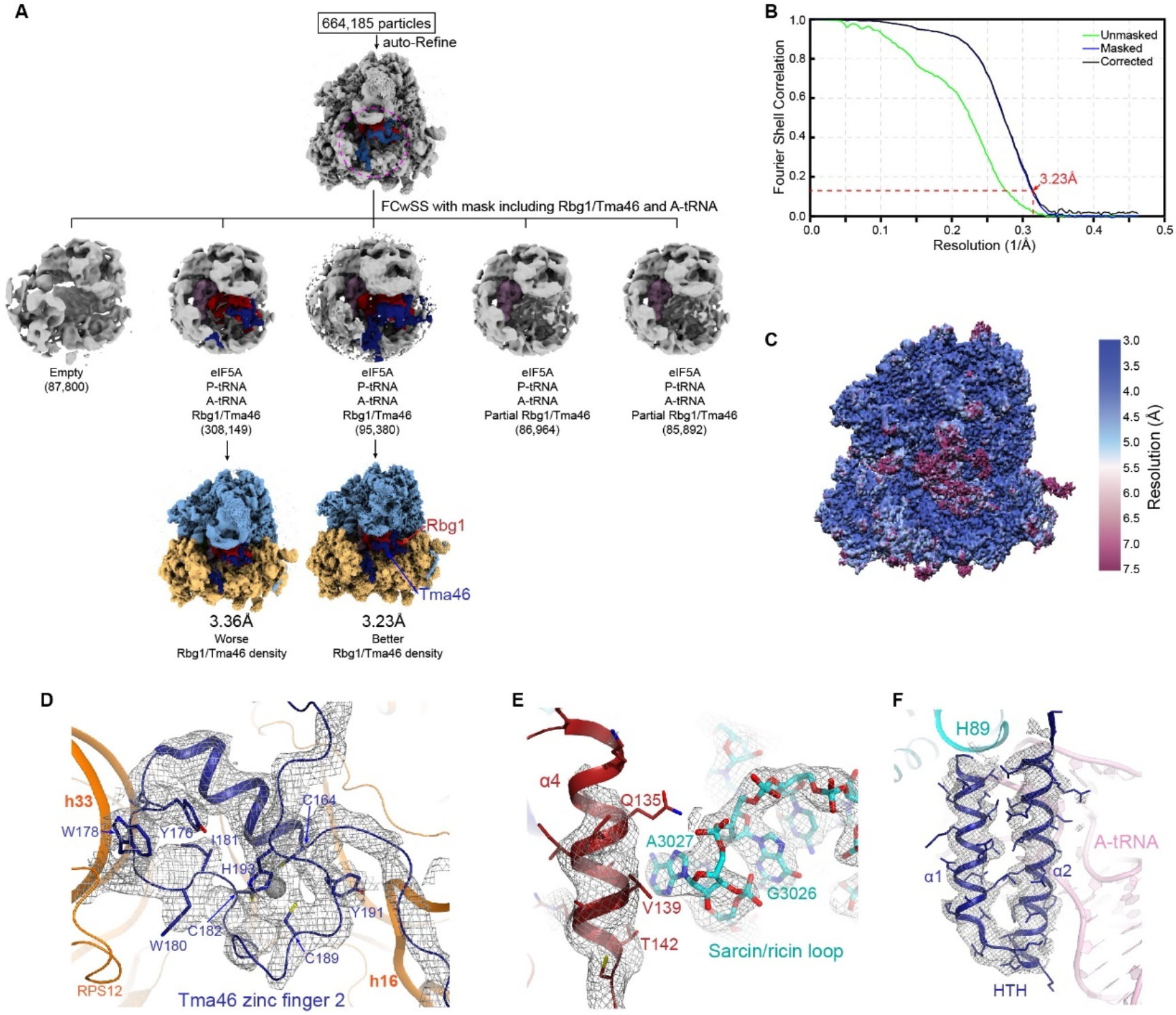
Structure determination of Rbg1/Tma46 on the ribosome and map quality. **A**. Focused classification with subtracted signal (FCwSS) with the mask over the Rbg1/Tma46 was performed on the selected 664,185 particles from 2D classification. Two of the five classes bound with Rbg1/Tma46 yielded 3.36 Å and 3.23 Å resolution, respectively. The two reconstructions are the same regarding to the conformation of proteins or ribosome. Only the 3.23 Å reconstruction which has better Rbg1/Tma46 density was used in this study. **B**. Gold-standard Fourier shell correlation (FSC) curves for the reported cryoEM reconstruction. **C**. The density map coloured by local resolution in surface view for the entire ribosomal complex. **D-F**. Representative electron microscopy maps showing the refined structures of zinc finger of Tma46 (D), sarcin/ricin loop of the large ribosomal subunit (E) and HTH domain of Rbg1 (F).

**Supplemental Figure 7.**
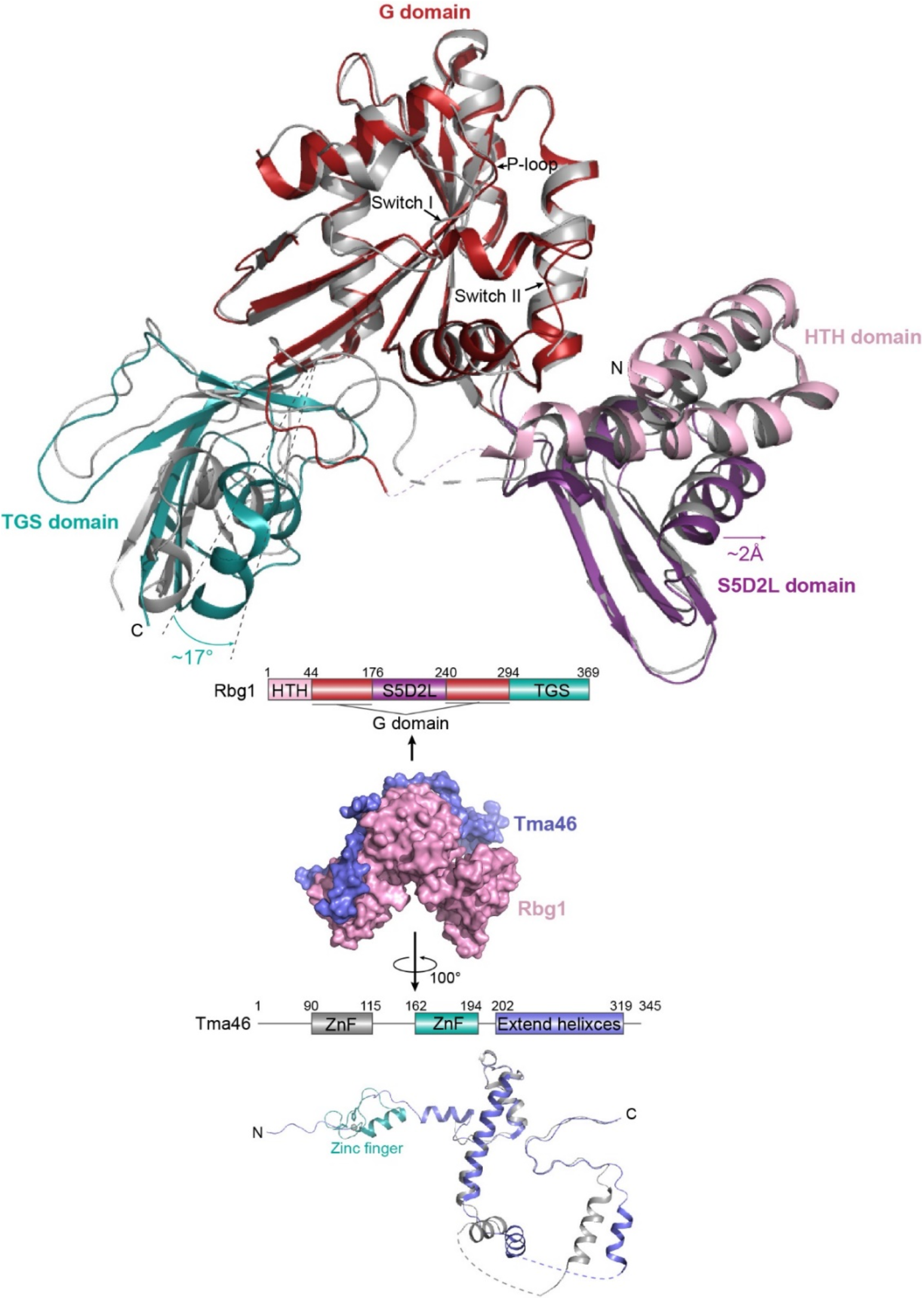
Conformation of Rbg1/Tma46 in the free form and ribosomal complex. Superposition of structures of Rbg1/Tma46 in the free form (PDB code: 4a9a, colored in grey) and bound to the ribosome (this study, multicolored) based on the alignment of the Rbg1 G-domain. When Rbg1/Tma46 binds to ribosome, the S5D2L domain extend out about 2Å and interacts with A-tRNA, the TGS domain rotate about 17° and interacts with h16 of 18S rRNA. The conformation of Tma46 changed due to the rotation of TGS domain in Rbg1.

**Supplemental Figure 8.**
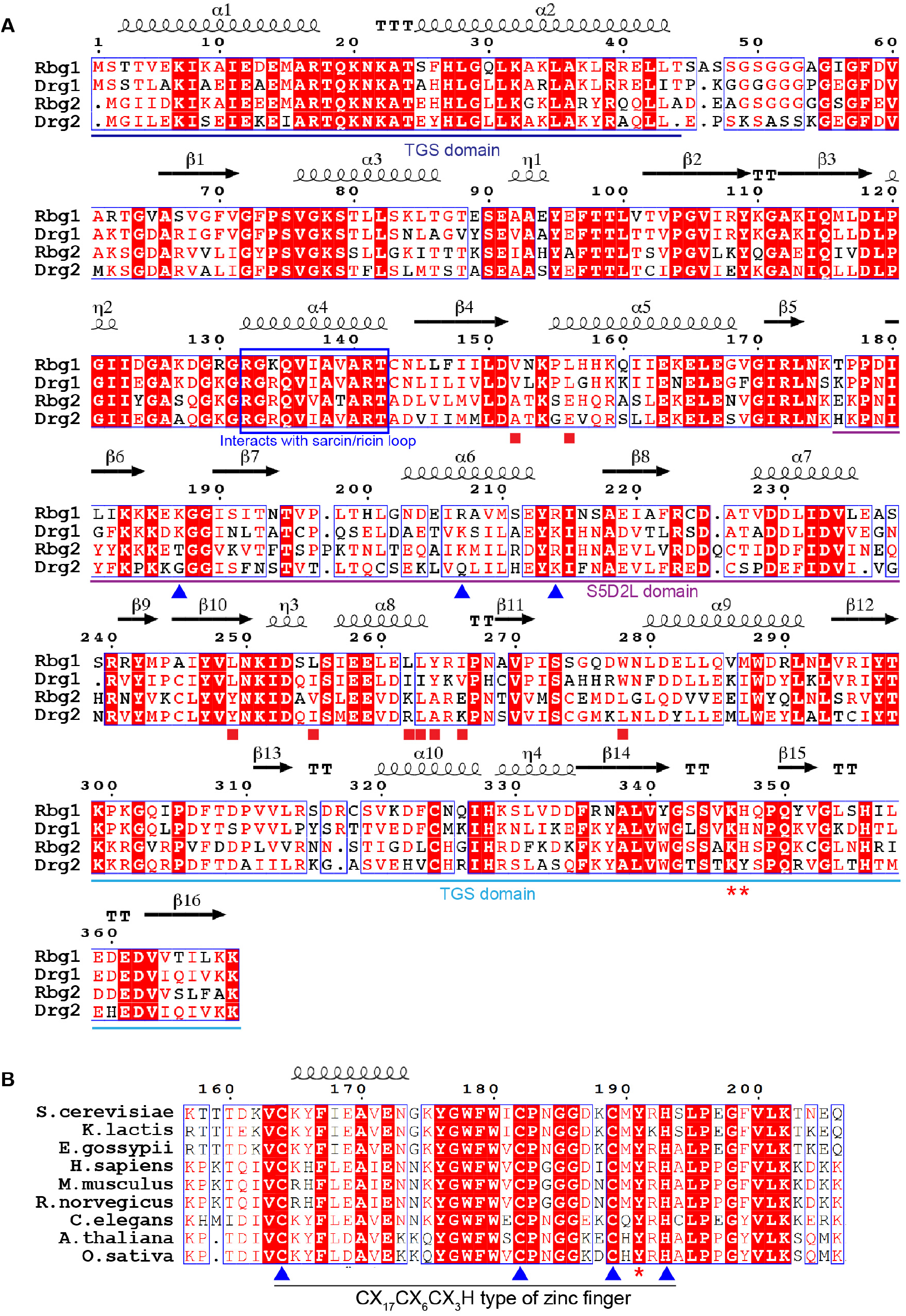
Sequence alignment of Rbg1 with Rbg2 and the zinc finger region of Tma46. **A**. Sequences of Rbg1 and Rbg2 in S. Cerevisiae, as well as Drg1 and Drg2 in Homo Sapiens Drg1, Drg2 were aligned. Secondary structures were shown base on the structure of Rbg1 obtained in this study. Rbg1 residues that interact with 18S rRNA and A-tRNA were indicated with the red star 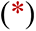 and blue triangle 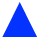, respectively. The region in Rbg1 helix α4 that interacts with the sarcin-ricin loop is highlighted with a blue box. Red squares 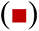 indicate residues in Rbg1 that interact with Tma46. **B**. Sequences of the second zinc finger of Tma46 in S. cerevisiae and homologues from other species were aligned. The CCCH motif was labeled in blue triangle 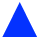 and the conserved tyrosine that stacks with A506 of 16S was labeled in red star 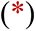.

## Supplementary Tables

**Supplementary Table 1.**
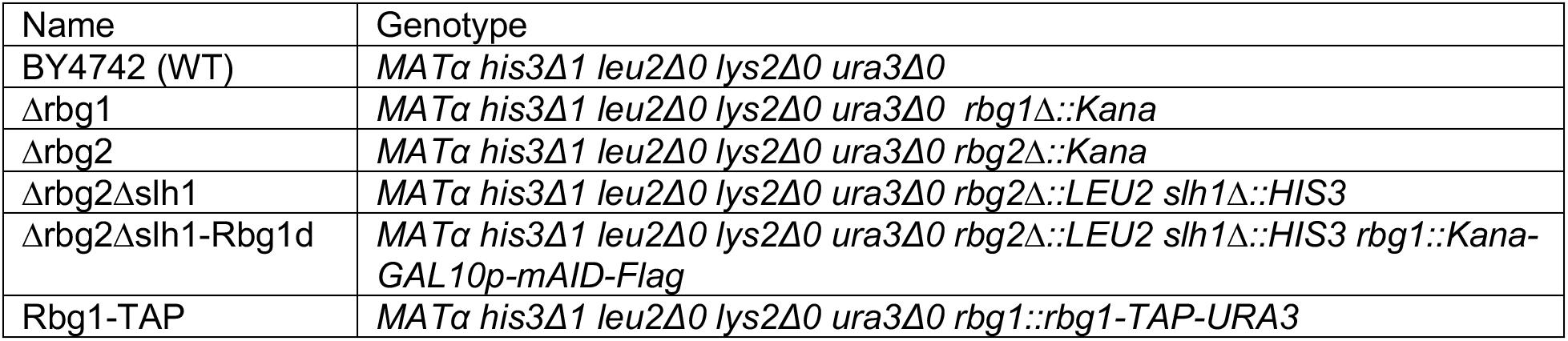
List of yeast strains used in the study

**Supplementary Table 2.** List of Plasmids used in the study

| Name | Details |
| --- | --- |
| pGAL10-mAID | Sequences including a KanMX selection marker, GAL10p, and mAID-3xFlag were inserted into the pET22b vector sequentially. Used to insert the mAID-3xFlag into the N-terminal of Rbg1 chromosomally. |
| p415G-TAP-URA3 | TAP tag, followed by a URA3 selection marker were inserted into the multiple cloning site of the p415 vector. Used to insert the TAP tag into the C-terminal of Rbg1 chromosomally. |
| pPGK1 | Empty vector modified from p415G, in which the GAL10 promoter was replaced by PGK1 promoter. |
| pPGK1-Rbg1 | Overexpress the wildtype Rbg1 using a PGK1 promoter in the Δrbg1 strain to restore the Rbg1 protein level. |
| pET28a-Rbg1 | Co-overexpress Rbg1/Tma46 in *E.coli* BL21(DE3) cells. |
| pET22b-Tma46 |  |
| pET22b-Tma46ΔZnF | Co-overexpress with pET28a-Rbg1 in *E.coli* BL21(DE3) cells to obtain the Rbg1/Tma46ΔZnF complex. |

**Supplementary Table 3.**
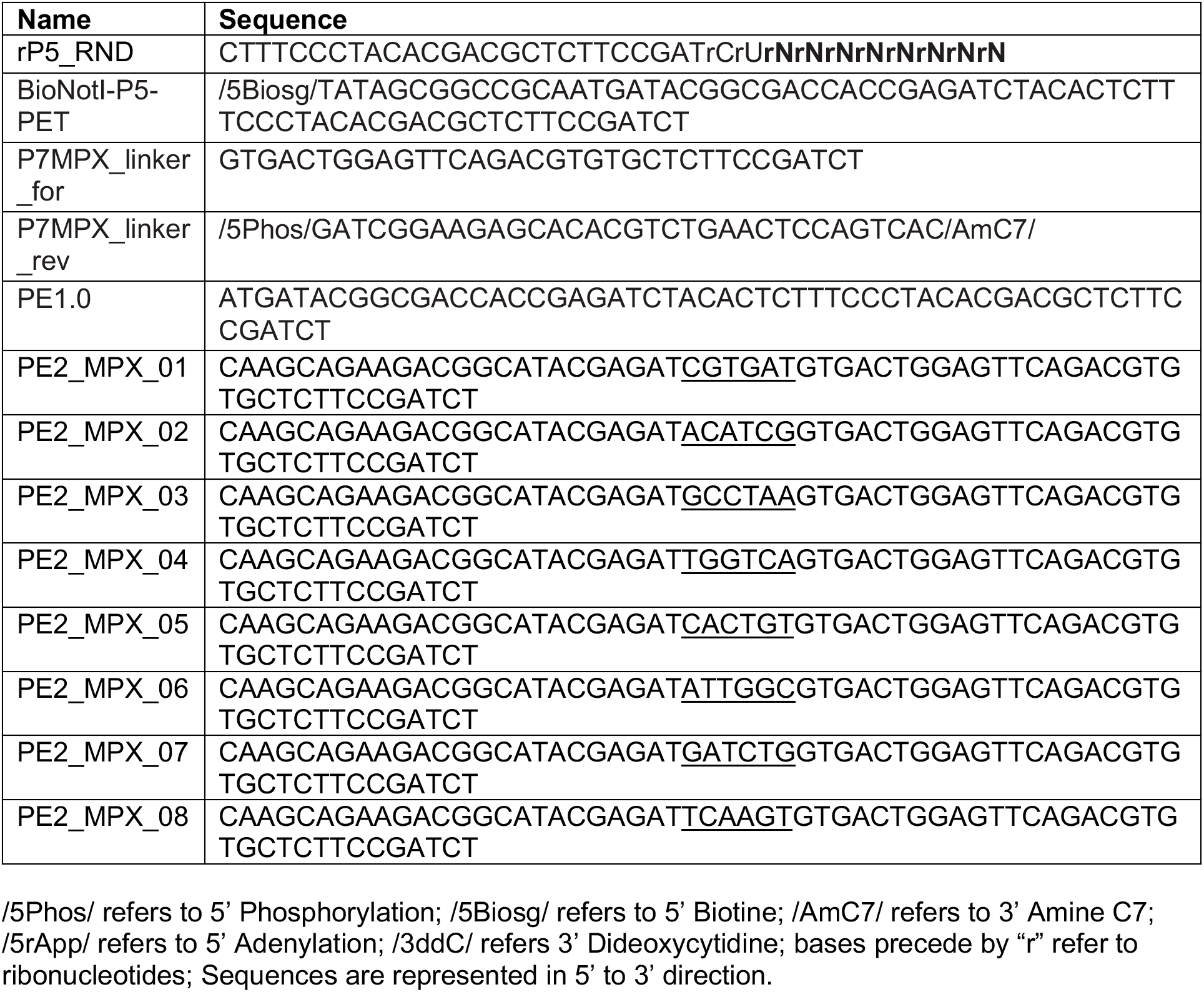
List of oligonucleotides used in the study

**Supplementary Table 4.** Data collection and model statistics

|  |  |
| --- | --- |
| **Data Collection** |  |
| Particles | 1,222,865 |
| Pixel size (Å) | 1.08 |
| Defocus range (μm) | -0.7 to -3.0 |
| Voltage (kV) | 300 |
| Electron dose (e<sup>-</sup>Å<sup>-2</sup>) | 30 |
| **Model composition** |  |
| Non-hydrogen atoms | 212,595 |
| Protein residues | 12,322 |
| RNA bases | 5,441 |
| Ligands (Zn<sup>2+</sup>/Mg<sup>2+</sup>) | 8/290 |
| **Refinement** |  |
| Resolution (Å) | 3.23 |
| Map sharpening B-factor (Å<sup>2</sup>) | -11 |
| FSC | 0.88 |
| **Rms deviations** |  |
| Bond lengths (Å) | 0.008 |
| Bond angles (°) | 0.807 |
| **Validation (proteins)** |  |
| Molprobity score | 2.12 |
| Clashscore, all atoms | 12.07 |
| Rotamer outliers (%) | 0.03 |
| **Ramachandran plot** |  |
| Favored (%) | 90.67 |
| Outlier (%) | 0.38 |
| **Validation (RNA)** |  |
| Correct sugar puckers (%) | 98.42% |
| Good backbone conformations (%) | 79.53% |

## References Cited

1. Balchin, D., Hayer-Hartl, M. & Hartl, F.U. In vivo aspects of protein folding and quality control. Science 353, aac4354 (2016).

2. Labbadia, J. & Morimoto, R.I. The biology of proteostasis in aging and disease. Annu Rev Biochem 84, 435–64 (2015).

3. Wolff, S., Weissman, J.S. & Dillin, A. Differential scales of protein quality control. Cell 157, 52–64 (2014).

4. . D’Orazio, K.N. & Green, R. Ribosome states signal RNA quality control. Mol Cell (2021).

5. Joazeiro, C.A.P. Mechanisms and functions of ribosome-associated protein quality control. Nat Rev Mol Cell Biol (2019).

6. Brandman, O. & Hegde, R.S. Ribosome-associated protein quality control. Nat Struct Mol Biol 23, 7–15 (2016).

7. Kumar, S., Tomooka, Y. & Noda, M. Identification of a set of genes with developmentally down-regulated expression in the mouse brain. Biochem Biophys Res Commun 185, 1155–61 (1992).

8. Sazuka, T. et al. Expression of DRG during murine embryonic development. Biochem Biophys Res Commun 189, 371–7 (1992).

9. Leipe, D.D., Wolf, Y.I., Koonin, E.V. & Aravind, L. Classification and evolution of P-loop GTPases and related ATPases. J Mol Biol 317, 41–72 (2002).

10. Sazuka, T., Tomooka, Y., Ikawa, Y., Noda, M. & Kumar, S. DRG: a novel developmentally regulated GTP-binding protein. Biochem Biophys Res Commun 189, 363–70 (1992).

11. Li, B. & Trueb, B. DRG represents a family of two closely related GTP-binding proteins. Biochim Biophys Acta 1491, 196–204 (2000).

12. Ishikawa, K., Azuma, S., Ikawa, S., Semba, K. & Inoue, J. Identification of DRG family regulatory proteins (DFRPs): specific regulation of DRG1 and DRG2. Genes Cells 10, 139–50 (2005).

13. Francis, S.M., Gas, M.E., Daugeron, M.C., Bravo, J. & Seraphin, B. Rbg1-Tma46 dimer structure reveals new functional domains and their role in polysome recruitment. Nucleic Acids Res 40, 11100–14 (2012).

14. O’Connell, A., Robin, G., Kobe, B. & Botella, J.R. Biochemical characterization of Arabidopsis developmentally regulated G-proteins (DRGs). Protein Expr Purif 67, 88–95 (2009).

15. Song, H. et al. Overexpression of DRG2 Increases G2/M Phase Cells and Decreases Sensitivity to Nocodazole-Induced Apoptosis. The Journal of Biochemistry 135, 331–335 (2004).

16. Mahajan, M.A., Park, S.T. & Sun, X.H. Association of a novel GTP binding protein, DRG, with TAL oncogenic proteins. Oncogene 12, 2343–50 (1996).

17. Zhao, X.-F. & Aplan, P.D. SCL binds the human homologue of DRG in vivo1The DRG cDNA sequence has been deposited in GenBank under accession number AF078103.1. Biochimica et Biophysica Acta (BBA) - Molecular Cell Research 1448, 109–114 (1998).

18. Schenker, T., Lach, C., Kessler, B., Calderara, S. & Trueb, B. A novel GTP-binding protein which is selectively repressed in SV40 transformed fibroblasts. J Biol Chem 269, 25447–53 (1994).

19. Daugeron, M.C., Prouteau, M., Lacroute, F. & Seraphin, B. The highly conserved eukaryotic DRG factors are required for efficient translation in a manner redundant with the putative RNA helicase Slh1. Nucleic Acids Res 39, 2221–33 (2011).

20. Pelechano, V., Wei, W. & Steinmetz, L.M. Widespread Co-translational RNA Decay Reveals Ribosome Dynamics. Cell 161, 1400–12 (2015).

21. Decourty, L. et al. Linking functionally related genes by sensitive and quantitative characterization of genetic interaction profiles. Proc Natl Acad Sci U S A 105, 5821–6 (2008).

22. Matsuo, Y. et al. Ubiquitination of stalled ribosome triggers ribosome-associated quality control. Nat Commun 8, 159 (2017).

23. Sitron, C.S., Park, J.H. & Brandman, O. Asc1, Hel2, and Slh1 couple translation arrest to nascent chain degradation. RNA 23, 798-810 (2017).

24. D’Orazio, K.N. et al. The endonuclease Cue2 cleaves mRNAs at stalled ribosomes during No Go Decay. Elife 8(2019).

25. Ikeuchi, K. et al. Collided ribosomes form a unique structural interface to induce Hel2- driven quality control pathways. EMBO J 38(2019).

26. Sugiyama, T. et al. Sequential Ubiquitination of Ribosomal Protein uS3 Triggers the Degradation of Non-functional 18S rRNA. Cell Rep 26, 3400–3415 e7 (2019).

27. Nishimura, K. & Kanemaki, M.T. Rapid Depletion of Budding Yeast Proteins via the Fusion of an Auxin-Inducible Degron (AID). Curr Protoc Cell Biol 64, 20 9 1–16 (2014).

28. Schuller, A.P., Wu, C.C., Dever, T.E., Buskirk, A.R. & Green, R. eIF5A Functions Globally in Translation Elongation and Termination. Mol Cell 66, 194–205 e5 (2017).

29. Guydosh, N.R. & Green, R. Dom34 rescues ribosomes in 3’ untranslated regions. Cell 156, 950–62 (2014).

30. Grollman, A.P. Inhibitors of protein biosynthesis. II. Mode of action of anisomycin. J Biol Chem 242, 3226–33 (1967).

31. Juszkiewicz, S. & Hegde, R.S. Initiation of Quality Control during Poly(A) Translation Requires Site-Specific Ribosome Ubiquitination. Mol Cell 65, 743–750.e4 (2017).

32. Gutierrez, E. et al. eIF5A promotes translation of polyproline motifs. Mol Cell 51, 35–45 (2013).

33. Pelechano, V. & Alepuz, P. eIF5A facilitates translation termination globally and promotes the elongation of many non polyproline-specific tripeptide sequences. Nucleic Acids Res 45, 7326–7338 (2017).

34. Doma, M.K. & Parker, R. Endonucleolytic cleavage of eukaryotic mRNAs with stalls in translation elongation. Nature 440, 561–564 (2006).

35. Letzring, D.P., Dean, K.M. & Grayhack, E.J. Control of translation efficiency in yeast by codon–anticodon interactions. RNA 16, 2516–2528 (2010).

36. Kuroha, K. et al. Receptor for activated C kinase 1 stimulates nascent polypeptide- dependent translation arrest. EMBO reports 11, 956–961 (2010).

37. Yan, L.L., Simms, C.L., McLoughlin, F., Vierstra, R.D. & Zaher, H.S. Oxidation and alkylation stresses activate ribosome-quality control. Nat Commun 10, 5611 (2019).

38. Vazquez-Laslop, N., Thum, C. & Mankin, A.S. Molecular mechanism of drug-dependent ribosome stalling. Mol Cell 30, 190–202 (2008).

39. Dimitrova, L.N., Kuroha, K., Tatematsu, T. & Inada, T. Nascent peptide-dependent translation arrest leads to Not4p-mediated protein degradation by the proteasome. J Biol Chem 284, 10343–52 (2009).

40. Wilson, D.N. & Beckmann, R. The ribosomal tunnel as a functional environment for nascent polypeptide folding and translational stalling. Curr Opin Struct Biol 21, 274–82 (2011).

41. Chandrasekaran, V. et al. Mechanism of ribosome stalling during translation of a poly(A) tail. Nat Struct Mol Biol 26, 1132–1140 (2019).

42. Davis, A.R., Gohara, D.W. & Yap, M.N. Sequence selectivity of macrolide-induced translational attenuation. Proc Natl Acad Sci U S A 111, 15379–84 (2014).

43. Pochopien, A.A. et al. Structure of Gcn1 bound to stalled and colliding 80S ribosomes. Proc Natl Acad Sci U S A 118(2021).

44. Wout, P.K., Sattlegger, E., Sullivan, S.M. & Maddock, J.R. Saccharomyces cerevisiae Rbg1 protein and its binding partner Gir2 interact on Polyribosomes with Gcn1. Eukaryot Cell 8, 1061–71 (2009).

45. Schmidt, C. et al. Structure of the hypusinylated eukaryotic translation factor eIF-5A bound to the ribosome. Nucleic Acids Res 44, 1944–51 (2016).

46. Melnikov, S. et al. Crystal Structure of Hypusine-Containing Translation Factor eIF5A Bound to a Rotated Eukaryotic Ribosome. Journal of molecular biology 428, 3570–3576 (2016).

47. Schmidt, C. et al. The cryo-EM structure of a ribosome–Ski2-Ski3-Ski8 helicase complex. Science 354, 1431–1433 (2016).

48. Buschauer, R. et al. The Ccr4-Not complex monitors the translating ribosome for codon optimality. Science 368(2020).

49. Lu, L., Lv, Y., Dong, J., Hu, S. & Peng, R. DRG1 is a potential oncogene in lung adenocarcinoma and promotes tumor progression via spindle checkpoint signaling regulation. Oncotarget 7, 72795–72806 (2016).

50. Schellhaus, A.K. et al. Developmentally Regulated GTP binding protein 1 (DRG1) controls microtubule dynamics. Sci Rep 7, 9996 (2017).

51. Bailey, T.L. et al. MEME SUITE: tools for motif discovery and searching. Nucleic Acids Res 37, W202–8 (2009).

## REFERENCES

1. Baudin, A., Ozier-Kalogeropoulos, O., Denouel, A., Lacroute, F. & Cullin, C. A simple and efficient method for direct gene deletion in Saccharomyces cerevisiae. Nucleic Acids Res 21, 3329–30 (1993).

2. Longtine, M.S. et al. Additional modules for versatile and economical PCR-based gene deletion and modification in Saccharomyces cerevisiae. Yeast 14, 953–61 (1998).

3. Daugeron, M.C., Prouteau, M., Lacroute, F. & Seraphin, B. The highly conserved eukaryotic DRG factors are required for efficient translation in a manner redundant with the putative RNA helicase Slh1. Nucleic Acids Res 39, 2221–33 (2011).

4. Schuller, A.P., Wu, C.C., Dever, T.E., Buskirk, A.R. & Green, R. eIF5A Functions Globally in Translation Elongation and Termination. Mol Cell 66, 194–205 e5 (2017).

5. Acker, M.G., Kolitz, S.E., Mitchell, S.F., Nanda, J.S. & Lorsch, J.R. Reconstitution of yeast translation initiation. Methods Enzymol 430, 111–45 (2007).

6. Francis, S.M., Gas, M.E., Daugeron, M.C., Bravo, J. & Seraphin, B. Rbg1-Tma46 dimer structure reveals new functional domains and their role in polysome recruitment. Nucleic Acids Res 40, 11100–14 (2012).

7. Xu, X., Lambrecht, A.D. & Xiao, W. Yeast survival and growth assays. Methods Mol Biol 1163, 183–91 (2014).

8. Zeng, F. & Jin, H. Peptide release promoted by methylated RF2 and ArfA in nonstop translation is achieved by an induced-fit mechanism. RNA 22, 49–60 (2016).

9. Pelechano, V., Wei, W. & Steinmetz, L.M. Genome-wide quantification of 5’- phosphorylated mRNA degradation intermediates for analysis of ribosome dynamics. Nat Protoc 11, 359–76 (2016).

10. Pelechano, V., Wei, W. & Steinmetz, L.M. Widespread Co-translational RNA Decay Reveals Ribosome Dynamics. Cell 161, 1400–12 (2015).

11. . Martin, M. Cutadapt removes adapter sequences from high-throughput sequencing reads. 2011 17, 3 (2011).

12. Langmead, B., Trapnell, C., Pop, M. & Salzberg, S.L. Ultrafast and memory-efficient alignment of short DNA sequences to the human genome. Genome Biol 10, R25 (2009).

13. Langmead, B. & Salzberg, S.L. Fast gapped-read alignment with Bowtie 2. Nat Methods 9, 357–9 (2012).

14. Bailey, T.L. et al. MEME SUITE: tools for motif discovery and searching. Nucleic Acids Res 37, W202–8 (2009).

15. Guydosh, N.R. & Green, R. Dom34 rescues ribosomes in 3’ untranslated regions. Cell 156, 950–62 (2014).

16. Zeng, F. et al. Structural basis of co-translational quality control by ArfA and RF2 bound to ribosome. Nature 541, 554–557 (2017).

17. Zheng, S.Q. et al. MotionCor2: anisotropic correction of beam-induced motion for improved cryo-electron microscopy. Nat Methods 14, 331–332 (2017).

18. Rohou, A. & Grigorieff, N. CTFFIND4: Fast and accurate defocus estimation from electron micrographs. J Struct Biol 192, 216–21 (2015).

19. Zivanov, J. et al. New tools for automated high-resolution cryo-EM structure determination in RELION-3. Elife 7(2018).

20. Scheres, S.H. & Chen, S. Prevention of overfitting in cryo-EM structure determination. Nat Methods 9, 853–4 (2012).

21. Ben-Shem, A. et al. The Structure of the Eukaryotic Ribosome at 3.0 Å Resolution. Science 334, 1524–1529 (2011).

22. Pettersen, E.F. et al. UCSF Chimera--a visualization system for exploratory research and analysis. J Comput Chem 25, 1605–12 (2004).

23. Emsley, P., Lohkamp, B., Scott, W.G. & Cowtan, K. Features and development of Coot. Acta Crystallographica Section D 66, 486–501 (2010).

24. Schmidt, C. et al. Structure of the hypusinylated eukaryotic translation factor eIF-5A bound to the ribosome. Nucleic Acids Res 44, 1944–51 (2016).

25. Waterhouse, A. et al. SWISS-MODEL: homology modelling of protein structures and complexes. Nucleic Acids Research 46, W296–W303 (2018).

26. Williams, C.J. et al. MolProbity: More and better reference data for improved all-atom structure validation. Protein Science 27, 293–315 (2018).

27. Schrodinger, LLC. The PyMOL Molecular Graphics System, Version 1.8. (2015).

## REFERENCES

1. Bailey, T.L. et al. MEME SUITE: tools for motif discovery and searching. Nucleic Acids Res 37, W202–8 (2009).

2. Huang da, W., Sherman, B.T. & Lempicki, R.A. Bioinformatics enrichment tools: paths toward the comprehensive functional analysis of large gene lists. Nucleic Acids Res 37, 1–13 (2009).

